# Bacterial Stress Responses Lower mRNA-Protein Level Correlations

**DOI:** 10.64898/2026.03.12.711437

**Authors:** Sena G. Süer, Jérôme Arnoux, Yi Y. Lim, Ganeshwari Dhurve, Rabia Şen, Cemal Erdem, André Mateus, Kemal Avican

## Abstract

Diverse bacterial pathogens have evolved complex regulatory mechanisms to adapt to various environmental stresses during infection. The uncertainty in mRNA-protein levels in response to environmental stressors complicates our understanding of bacterial physiology and their adaptation to stressful environments. To examine this issue, we have integrated transcriptomics and proteomics data on three human bacterial pathogens *Salmonella enterica* Typhimurium, *Yersinia pseudotuberculosis*, and *Staphylococcus aureus* under ten infection-relevant stress conditions. We observed positive correlations between mRNA and protein levels, which were decreased under different stress conditions. Essential genes exhibited higher expression levels with lower variation across the conditions and stronger mRNA-protein correlations compared to non-essential genes, highlighting their critical role in bacterial adaptability and survival. Moreover, we identified a substantial number of genes with stress-induced non-correlating mRNA-protein levels, especially under conditions triggering strong stress responses. Particularly this level was dramatically lowered for osmotic stress specific genes affected by impaired translational activity under osmotic stress. Our findings highlight the prevalence of non-correlating mRNA-protein levels and the potential role of post-translational modifications in modulating protein levels in response to environmental stressors during infection. This study provides a comprehensive framework for integrating transcriptomics and proteomics data and identifies potential gene products that might significantly impact the ability of diverse bacterial pathogens to adapt to hostile infection environments.

**Significance Statement:** Understanding how bacteria adapt to host environments is crucial for combating infections. We employed an integrative transcriptomics and proteomics approach to investigate mRNA-protein correlations in three clinically relevant pathogens under ten infection-relevant stress conditions. We identified genes whose mRNA-protein relationships are significantly disrupted by specific stressors. This study provides a deeper understanding of mRNA-protein level uncertainty, coupling it to stress responses known to trigger critical post-transcriptional and post-translational regulations. These findings help reveal novel mechanisms of bacterial adaptation and pathogenesis. By providing a comprehensive, multi-species dataset, this study serves as a foundational resource for infection biology, driving future advancements in understanding complex regulatory networks and identifying potential antimicrobial targets.

## Introduction

Bacterial pathogens have evolved complex mechanisms to adapt to different environmental stresses in the host such as nutrient limitations, temperature or pH shifts, and antimicrobial challenges during infection. They respond to deleterious effects of such stressors via stress response mechanisms, involving complex gene regulatory networks, for survival. Therefore, comprehensive understanding of bacterial stress response mechanisms is critical for development of novel therapeutic approaches to resolve bacterial infections. One useful approach is through investigation of global transcriptomes of diverse bacterial pathogens under a variety of stress conditions that mimic host environments. This provides information on regulation across diverse stress conditions that could be used in attempts to reveal complex gene regulations and identify key stress response genes. We previously performed a large-scale transcriptomic study of 32 human bacterial pathogens, PATHOgenex RNA Atlas (1), aiming to understand bacterial stress responses under 11 *in vivo* mimicking stress conditions. The study revealed a dynamic regulation of gene expression under different stress conditions and suggested a set of genes that has potential to be candidate as new antimicrobial targets. Co-PATHOgenex (2), the continuation of the PATHOgenex study, identifies stress-specific stimulons by generating gene co-expression modules. These stimulons represent sets of genes with unique expression patterns under specific stressors, providing insights into the stress adaptation to different stressors.

Yet the main functional units shaping the cellular response are proteins and there is, generally, a weak correlation between mRNA and protein levels (3–11). This highlights the complex nature of gene regulation and the diverse post-transcriptional factors affecting protein stability, and abundance (12). It is known that transcription and translation could be uncoupled constitutively or under certain conditions, such as stress conditions, at a global or at a gene-specific level (13, 14) Therefore, the relationship between mRNA and protein levels can vary depending on the conditions, and this variability has been the subject of extensive research and ongoing debate (12, 14–16). Nevertheless, mRNA levels have been shown to be positively correlated with protein levels in many different organisms (8, 17–25). Linking protein expression to gene expression can help to identify key proteins in response to specific stimuli or conditions and highlight mechanisms that drive bacterial adaptation, evasion of the immune system, and disease development (26–30). This is crucial for developing more effective diagnostic tools, treatment strategies, and vaccines for bacterial infections (31, 32). Therefore, integrated transcriptomic and proteomic studies have emerged as powerful tools for understanding the complex molecular mechanisms underlying bacterial pathogenesis (28). Despite the numerous studies integrating transcriptome and proteome data, efforts to identify genes where mRNA and protein levels *do not* correlate are relatively limited (17, 33). Focusing on such genes could offer valuable insight into the complex post-transcriptional and post-translational regulatory mechanisms that escape detection by analyses addressing correlated gene-expression patterns.

In this study, we employed an integrative transcriptomic and proteomic analysis of *Salmonella enterica* Typhimurium, *Yersinia pseudotuberculosis*, and *Staphylococcus aureus* under ten infection-relevant stress conditions to investigate mRNA-protein level correlations. We revealed that mRNA-protein correlations significantly varied under different stress conditions for the three species. Essential genes consistently showed higher mRNA and protein expression levels and stronger correlations, emphasizing their importance in bacterial survival. Stress conditions such as hypoxia and nutritional downshift exhibited the lowest correlations, suggesting that post-transcriptional and post-translational regulation play a significant role under these conditions. We identified plentiful genes that showed non-correlating mRNA-protein levels in the three species, as well as sets of genes that were species specific, particularly under conditions eliciting strong stress responses. Among all conditions tested, osmotic stress stood out as exhibiting the most pronounced mRNA-protein decoupling. Using the MOBILE (34) framework, we characterized the osmotic stress stimulon and linked the observed mRNA-protein decoupling to a specific impairment of translation initiation.

## Results

### Transcriptomic and proteomic analysis of three human bacterial pathogens under ten infection relevant stress conditions

We analyzed transcriptomic and proteomic datasets of three human bacterial pathogens including Gram-negative *S. enterica* Typhimurium SL1344, *Y. pseudotuberculosis* YPIII and Gram-positive *S. aureus* MSSA476. We used transcriptomes of the three pathogens under 10 infection relevant stress conditions together with unexposed control condition from PATHOgenex RNA Atlas (1) and complemented it with mass-spectrometry proteomic profiling under the same stressors (**Figure 1A**). As mass spectrometry-based proteomics often yields a lower coverage than RNA-seq, we compare detected transcripts and proteins across all conditions in three pathogens, focusing on protein coding sequences (CDSs) in the analysis. While almost all CDSs were detected by RNA-seq for the three pathogens (98% for *S. enterica* and *Y. pseudotuberculosis*, and 90% for *S. aureus*), proteomics could detect between 48% of the CDSs for *S. enterica* Typhimurium and *Y. pseudotuberculosis*, and 32% for *S. aureus* (**Figure 1B**). Detected proteins covered nearly the full spectrum of mRNA expressions, with a bias towards highly expressed mRNAs across the conditions for the three pathogens (**Figure 1C**).

**Figure 1.**
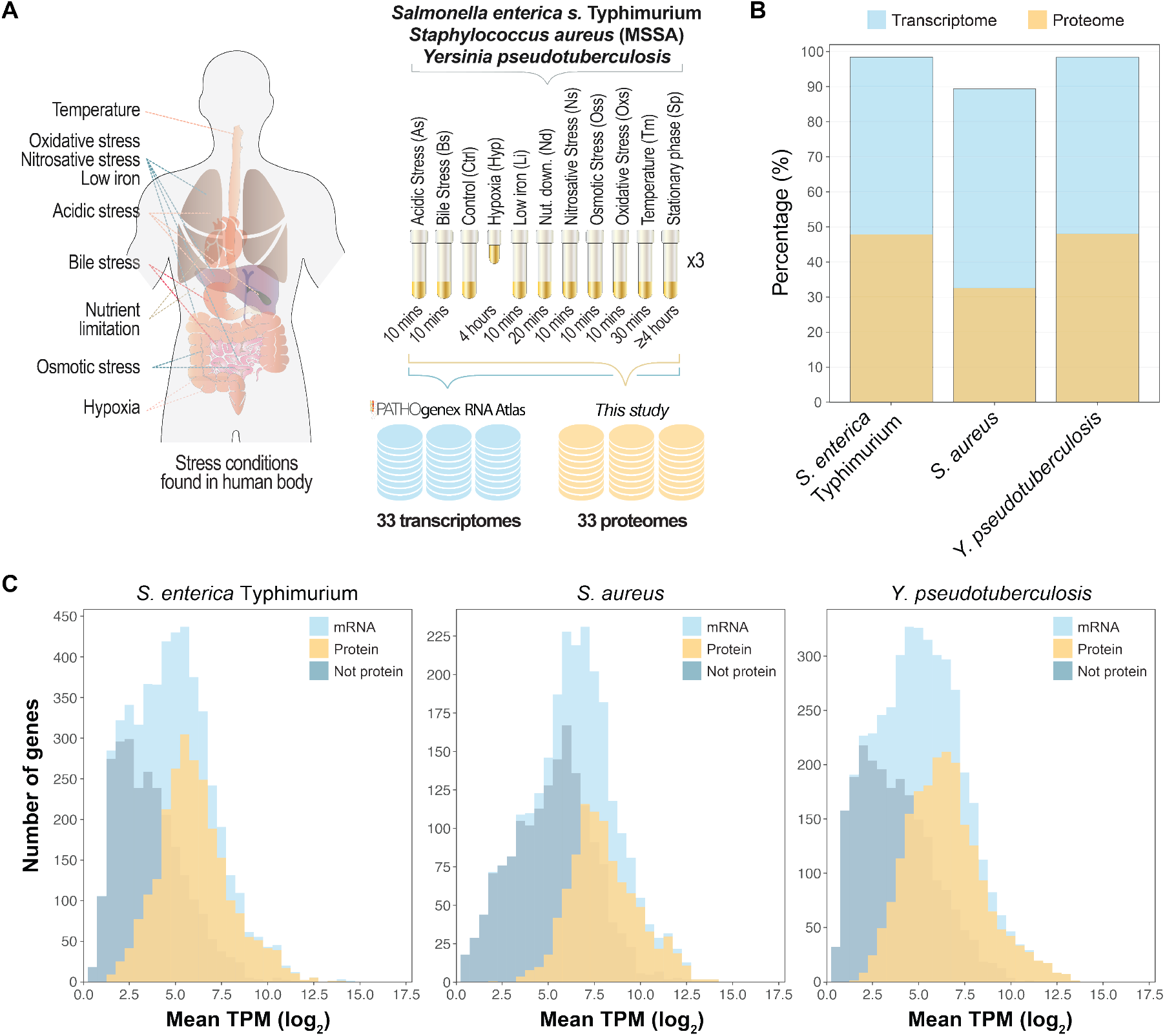
Transcriptomic and proteomic analysis of three human bacterial pathogens under ten infection relevant stress conditions. **(A)** Schematic presentation of data collection and data generation for gene expression on mRNA and protein levels under stress conditions. **(B)** Percentages of CDSs detected by transcriptomics and proteomics for the three species. **(C)** Distribution of transcript abundance across all conditions (light blue) in the three species; yellow indicates the fraction of proteins detected, while dark blue represents the transcripts for which no corresponding protein was detected.

### mRNA-protein level correlation under stress

The correlation between mRNA and protein levels has been previously explored in bacteria under individual growth conditions (18, 33, 35, 36). Our data allows us to assess whether mRNA levels can serve as an accurate indicator of protein levels in three human bacterial pathogens. To evaluate how different stressors affect the mRNA-protein level correlation, we calculated the intensity-Based Absolute Quantification (iBAQ) (15) **(Supplementary Table 1)** values for protein levels and compared to mRNA levels in Transcript Per Million (TPM) (37) for every stress condition including the unexposed control condition (1). We have previously shown that the three biological replicates of the RNA-seq samples clustered together (1). For the proteome dataset generated in this study, we conducted pairwise comparisons of the three replicates of each condition and computed Pearson correlation coefficient values (R-value). The R-value for pairwise comparison within the replicates of the same condition varied from 0.89-0.96 for the three pathogens **(Supplementary Figures 1-3)** showing strong similarities between the replicates. We then measured the correlation between mRNA and protein levels by calculating Pearson correlation coefficient values of the log2 transformed mean TPM and log2 transformed mean iBAQ of genes for each condition and pathogens **(Figure 2A, B, and C)**. For these measurements, we limited the analysis to the genes that are detected as mRNA and protein across all tested conditions. We observed that correlations between mRNA and protein levels under 11 conditions ranged from 0.48-0.68 for *S. enterica* Typhimurium, 0.54-0.72 for *S. aureus* and 0.59-0.76 for *Y. pseudotuberculosis* **(Figure 2A, B, and C)**. These correlation coefficients are comparable to those reported in previous studies across diverse biological systems (19, 20, 33, 36, 38). The variation in mRNA–protein correlation across stress conditions suggests that distinct molecular mechanisms, activated by each stressor, might modulate translational or post-translational regulations leading to differences.

**Table 1.** Classification of differentially expressed genes based on mRNA and protein regulation patterns.

| Cluster | mRNA regulation | Protein regulation | Description |
| --- | --- | --- | --- |
| 1 | ▼ Downregulated | ● Not changed | mRNA is downregulated in at least one stress condition, while protein level remains unchanged |
| 2 | ▼ Downregulated | ▲ Upregulated | mRNA is downregulated in at least one condition, while protein level is upregulated |
| 3 | ● Not changed | ▼ Downregulated | mRNA level remains unchanged across all conditions, while protein is downregulated |
| 4 | ● Not changed | ▲ Upregulated | mRNA level remains unchanged across all conditions, while protein is upregulated in at least one condition |
| 5 | ▲ Upregulated | ▼ Downregulated | mRNA is upregulated in at least one condition, while protein level is downregulated |
| 6 | ▲ Upregulated | ● Not changed | mRNA is upregulated in at least one condition, while protein level remains unchanged |

**Figure 2.**
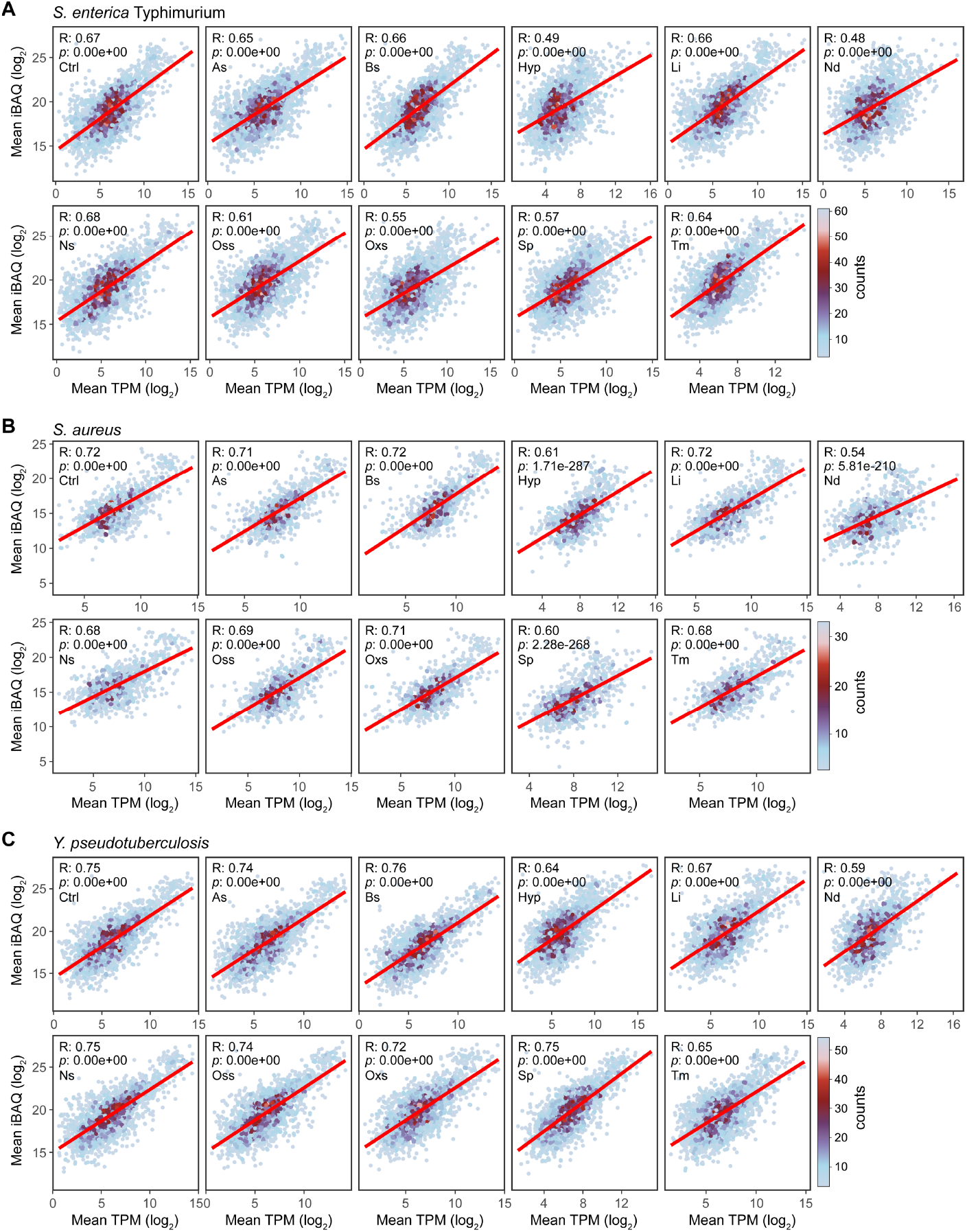
mRNA-protein levels show differing degrees of statistically significant positive correlation under different stress conditions and control conditions in **(A)** *S. enterica* Typhimurium, **(B)** *S. aureus* and **(C)** *Y. pseudotuberculosis*. mRNA and protein expression levels were taken as log_2_ transformed TPM and iBAQ values, respectively. The expression values of genes that are detected in all conditions as mRNA and protein were used in this analysis. Pearson correlation coefficient (R) shows positive correlation under all conditions in the three species. *p* values were calculated using Student’s t-test.

### Essential genes have higher mRNA-protein level correlations in the three species

The positive correlation between mRNA-protein levels for all detected genes prompted us to investigate subsets of genes with different correlations levels. Essential genes have been shown to have stronger correlations in comparison to non-essentials for *Pseudomonas aeruginosa* during different growth phases (17). Therefore, we sought to investigate mRNA-protein level correlation profiles of essential genes in the three species. Of the 354, 341, and 445 previously identified essential genes for *S. enterica* Typhimurium (39) *S. aureus* (40) and *Y. pseudotuberculosis* (41) respectively, we detected 300 (85%), 280 (82%), and 335 (75%) essential genes in our transcriptome and proteome datasets. We found that mean expression levels of mRNAs and proteins were significantly higher for essential genes in comparison to non-essentials for the three species (**Figure 3A and B)**. This was true for the expression levels in every condition **(Supplementary Figures 4 and 5**). Moreover, we observed that essential genes have higher mRNA-protein level correlation than all detected genes for all three species (**Figure 3C**), except for *S. enterica* Typhimurium and *Y. pseudotuberculosis* under nutritional downshift (Nd) stress condition. This distinct uncoupling in the Gram-negative species hints at divergent regulatory priorities during sudden starvation. As previously observed in other species (42, 43), essential genes showed significantly lower variance on both mRNA and protein expression across the 11 conditions for the three species (**Figure 3D and E**). These findings show that essential genes of *S. enterica* Typhimurium, *S. aureus*, and *Y. pseudotuberculosis* have high and stable mRNA and protein expression under a wide range of stress conditions with robust mRNA-protein level correlations.

**Figure 3.**
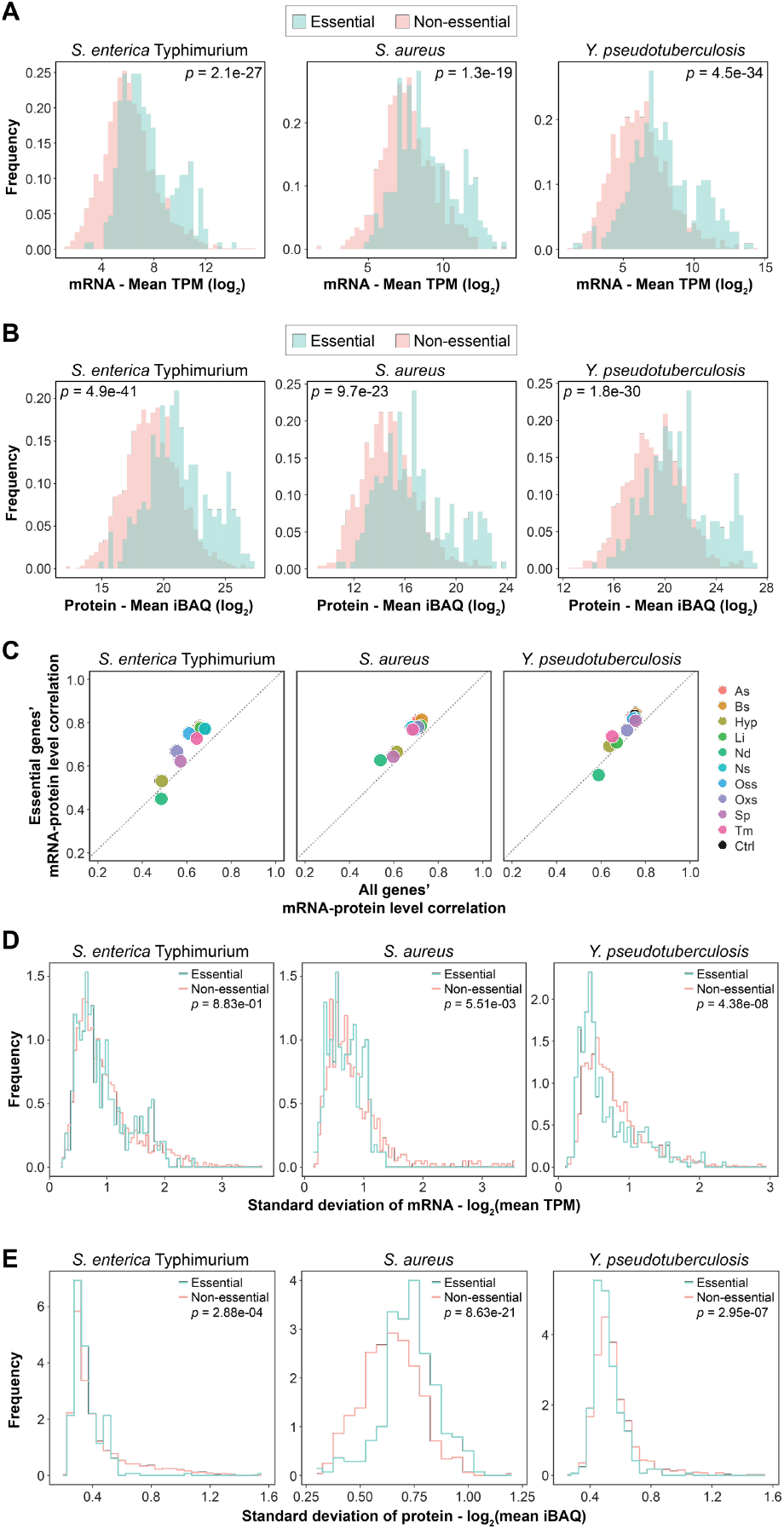
Essential genes of the three pathogens have higher mRNA and protein expressions with higher correlation and lower variance across the 11 conditions in comparison to all genes. **(A)** Mean mRNA expression frequency of essential and non-essential genes in the 11 conditions. **(B)** Mean protein expression frequency of essential and non-essential genes in the 11 conditions. **(C)** Essential genes and all genes mRNA-protein level correlation under 11 conditions for the three species. **(D)** Frequency of standard deviation of mRNA expression under 11 conditions for the three species. **(E)** Frequency of standard deviation of protein expression under 11 conditions for the three species. mRNA-protein level correlations were measured with Pearson correlation coefficient (R). *p* values were calculated using a Wilcoxon rank sum test.

**Figure 4.**
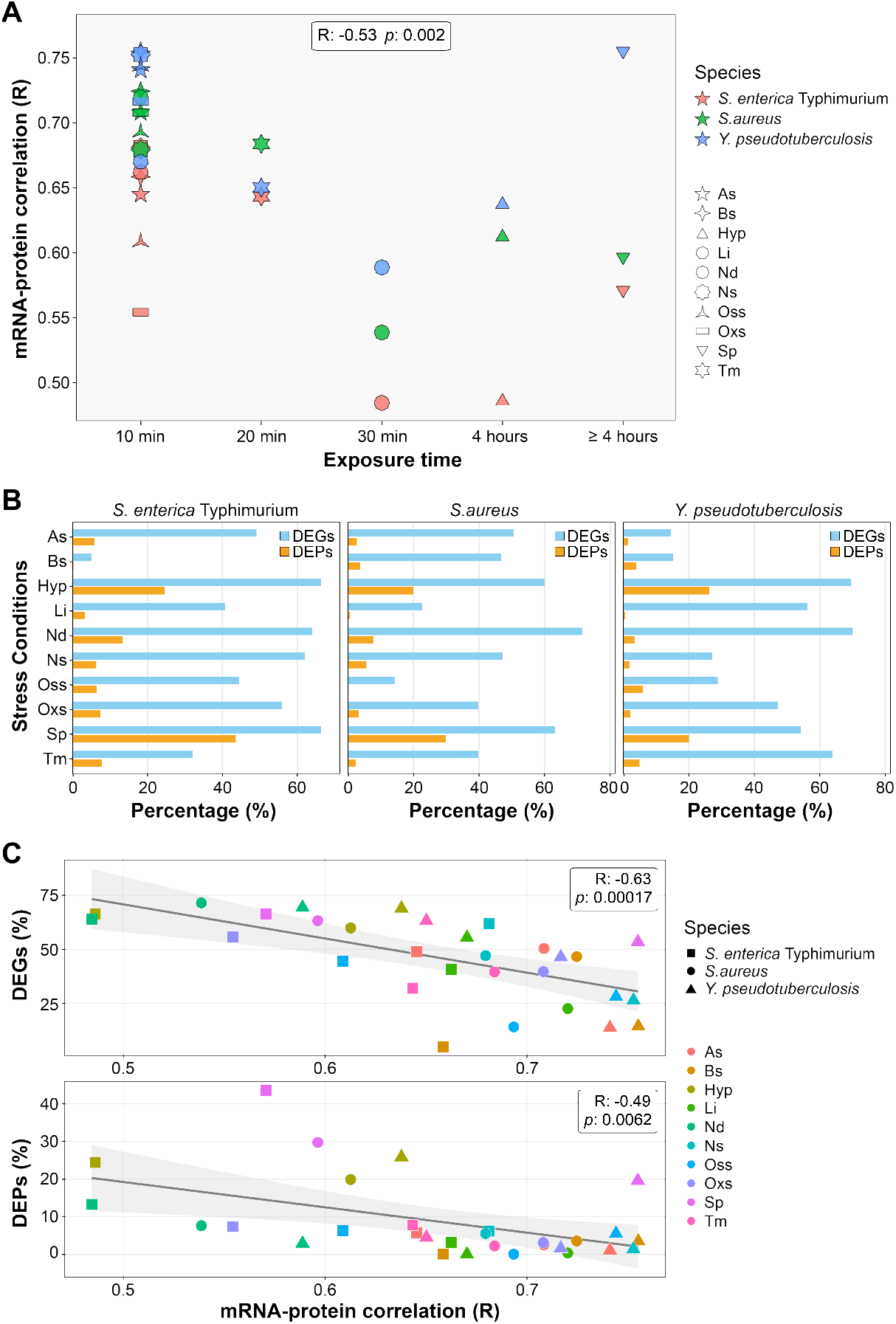
mRNA-protein level correlation lowers with increasing degree of stress response. **(A)** mRNA-protein level correlations for conditions exposed to stressors for 10 minutes, 20 minutes, 30 minutes, 4 hours, and longer than 4 hours in the three species. **(B)** Percentages of differentially expressed genes (DEGs) and proteins (DEPs) under the 10 stress conditions in comparison to unexposed control conditions for the three species. **(C)** Scatter plots showing negative correlation between percentages of DEGs (top) and DEPs (bottom) and mRNA-protein level correlation. Correlations were measured with Spearman correlation (R) for **(A)** and Pearson correlation (R) for **(C).** *p* values were calculated using Student’s t-test.

**Figure 5.**
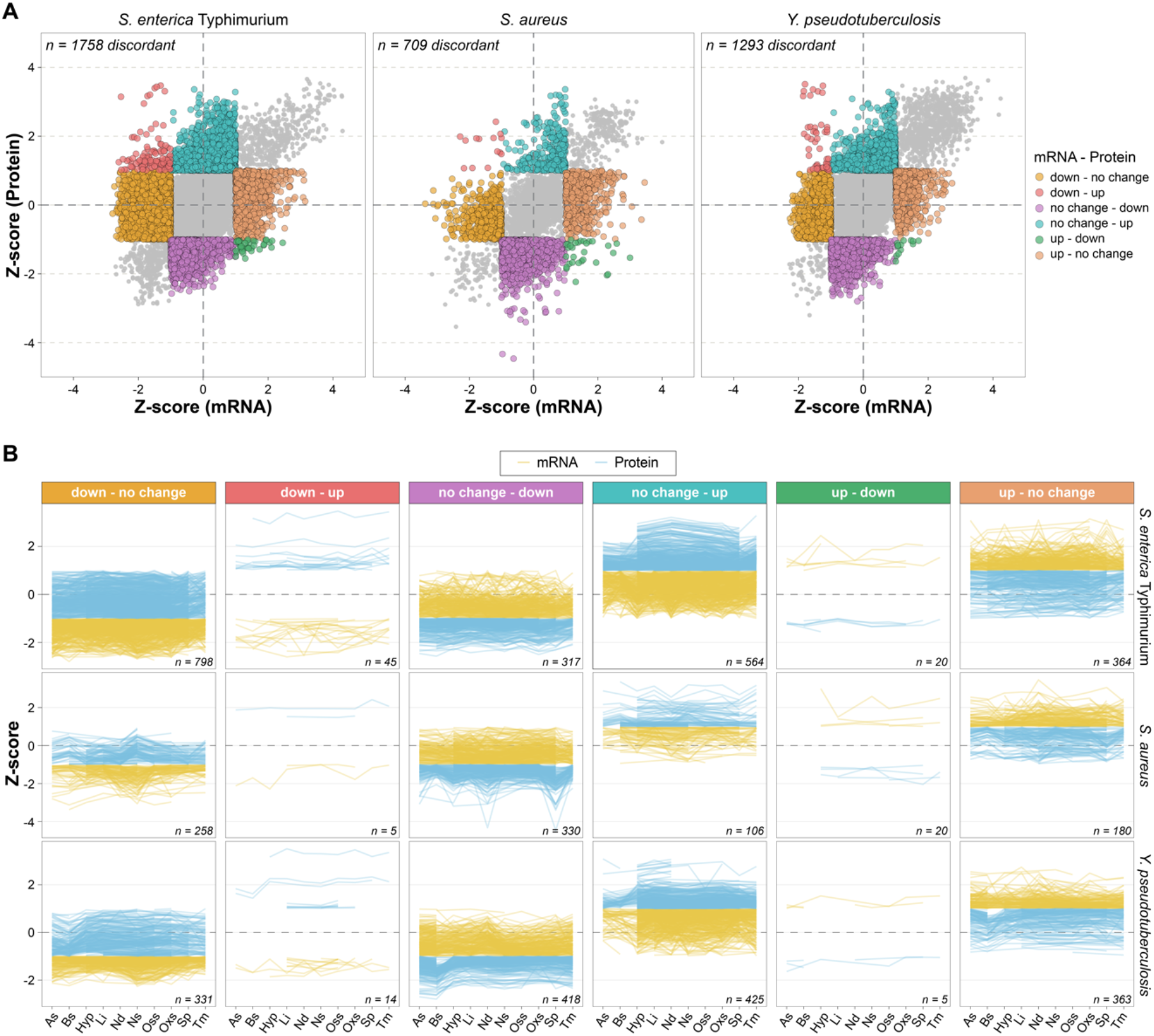
Stress response induces different patterns of discordant mRNA and protein level regulation. **(A)** Scatter plots with Z-scores of mRNA and protein levels for each of the 10 stress conditions in comparison to control for the three species. The 6 gene clusters whose mRNA and protein level have different discordant regulation patterns are shown in different colors. **(B)** Z-score patterns of mRNA and protein levels across 10 stress conditions for the 6 discordantly regulated gene clusters in each species. The number of discordantly regulated genes per cluster is indicated with “*n*”.

### Stress responses differentially affect mRNA-protein correlation

The variation in correlation under different stress conditions prompted us to investigate possible underlaying reasons. The mRNA-protein levels in Hypoxia (Hyp) and Nd showed the lowest correlation across three species **(Figure 2A, B, and C)**. Those stress conditions differ from others in the length of the exposure time: 4 hours for Hyp and 30 minutes for Nd, against 10 minutes for others **(Figure 1A)** besides condition itself. To determine whether exposure duration influences mRNA-protein level correlation, we examined the relationship between correlation and exposure time across all conditions. We observed a decline in mRNA-protein level correlation associated with longer exposure times (R = -0.53; *p* = 0.002**; Figure 4A)**. However, this effect may not be solely dependent on time; robust, stressor-specific responses triggered by conditions like Nd, stationary phase (Sp), and Temperature (Tm) likely act as confounding factors that drive this decrease.

To assess whether the degree of stress responses through differentially regulated mRNAs and proteins has an impact on mRNA-protein level correlations, we used differentially expressed genes (DEGs) and differentially expressed proteins (DEPs) for each condition in the three species by using unexposed samples as control. Before identifying DEPs, we evaluated 8 normalization methods from NormalyzerDE (44), on the iBAQ protein abundance data and found that Variance Stabilizing Normalization (VSN) was the most effective in clustering the condition together (**Supplementary Figure 6**). Then we used limma (45) differential expression package for the identification of DEPs with VSN normalized data. For DEGs, we used previously reported differential expression analysis of PATHOgenex RNA Atlas (1). We employed a *p*-value < 0.05 and a log2-Fold Change > |0.5| as thresholds to identify DEGs and DEPs for each condition. Notably, the number of DEGs were significantly higher compared to DEPs in all conditions for the three species **(Figure 4B)**. This can be linked to the stronger regulation of mRNA levels than of protein levels as previously observed in humans (16, 46). Moreover, percentages of DEGs and DEPs were highest under Hyp, Nd, and Sp conditions for the three pathogens (**Figure 4B**). Next, we tested whether the differential expressions have an impact on mRNA-protein level correlations. To do this, we measured the association between the percentages of DEGs and DEPs and the mRNA-protein level correlations with Pearson correlation coefficient. Our results revealed a strong negative correlation between differential expressions and mRNA-protein level correlation (**Figure 4C**). This suggests that other factors, such as post-transcriptional and post-translational regulation triggered by different stressors, might be influencing protein levels alongside the mRNA expression.

**Figure 6.**
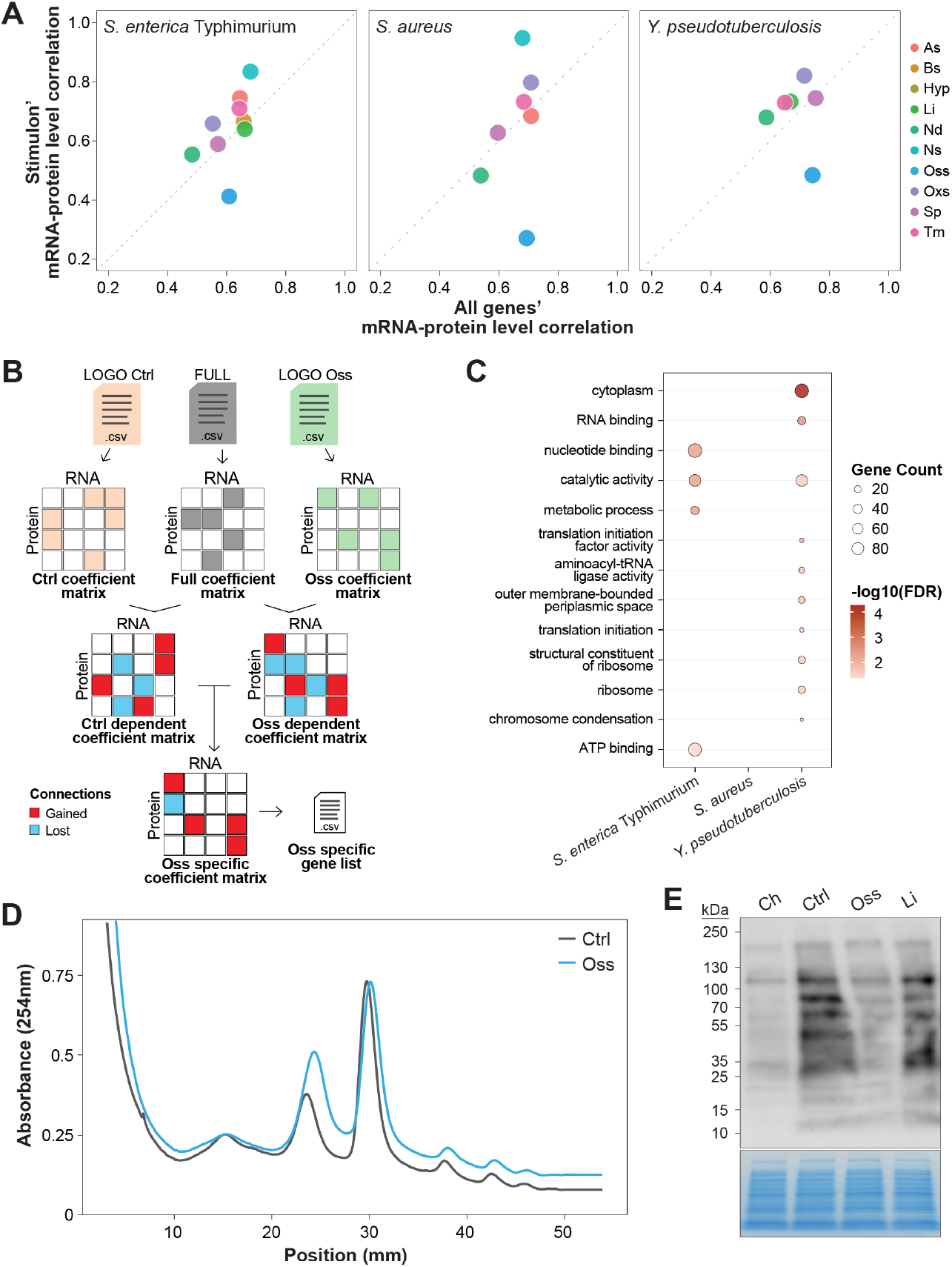
Impaired translation efficiency lowers mRNA-protein levels under osmotic stress. A) Stress specific stimulon genes and all genes mRNA-protein level correlation under 10 conditions for the three species. B) Schematic representation of MOBILE workflow for identifying Oss-specific mRNA-protein associations. The diagram outlines MOBILE analysis applied to transcriptomics and proteomics datasets from 11 conditions. Initial data from the control condition (LOGO Ctrl), the complete dataset (FULL), and the Oss condition (LOGO Oss) are used to construct corresponding base coefficient matrices. The FULL matrix is then compared against the Ctrl and Oss matrices to generate condition-dependent coefficient matrices, which highlight specific mRNA-protein connections that are either gained (red squares) or lost (blue squares). Finally, a comparison between the Ctrl-dependent and Oss-dependent matrices isolates the Oss-specific coefficient matrix, resulting in a targeted Oss-specific connections gene list. C) Over representation analysis (ORA) of Oss specific connections across three species, representing enrichment across biological process, molecular function, and cellular component. D) Polysome profiles of wt *Y. pseudotuberculosis* cells under control (Ctrl) and osmotic stress (0.5M NaCl) conditions. E) Western blot membrane (upper panel) and Coomassie blue-stained gel (lower panel) of total proteins extracted from wt *Y. pseudotuberculosis* cells under negative control (Ch), positive control (Ctrl), osmotic stress (Oss), and low iron (Li). Active translation of nascent proteins was detected using a monoclonal anti-puromycin antibody.

### Identification of genes whose mRNA-protein levels are discordantly regulated under stress conditions

The decreased correlation between mRNA-protein levels with increased number of DEGs and DEPs under specific stress conditions led us to investigate the genes whose mRNA-protein levels are discordantly regulated when the bacteria are exposed to stress conditions. We clustered genes into 6 groups (Error! Reference source not found.Error! Reference source not found.) corresponding to distinct combinations of differential expression patterns at the mRNA and protein levels in at least one stress condition (**Figure 5A and B and Supplementary Table 2**). In agreement with lowest correlation between mRNA-protein levels observed under Hyp, Nd, and Sp conditions (**Figure 2A-C**), discordantly regulated genes were mostly found under those conditions for the three species (**Supplementary Figure 7**). Noteworthy, Clusters 1 and 6, where mRNAs were regulated but not proteins, had the highest number of discordantly regulated genes. This is in accordance with higher number of differentially regulated mRNAs (DEGs) compared to differentially regulated proteins (DEPs) in all conditions for the three species (**Figure 4B**).

### Discordantly regulated genes are enriched in translation

To investigate whether the genes exhibiting discordant mRNA and protein levels under stress conditions are enriched in certain feature, we performed Gene Ontology (GO) (47) enrichment analysis. Interestingly, we found that biological processes such as translation, RNA processing, RNA modification, phosphorylation, and cell division, among other biological processes were commonly enriched for the three species, particularly when mRNA levels were changed but there was no change at the protein level (**Supplementary Figure 8 and Supplementary Table 3**). This suggests that genes encoding subunits of translation machinery together with other components of translation such as tRNA and rRNA were discordantly regulated under stress conditions for the three pathogens. In detail, proteins in Cluster 1 have stable levels despite mRNA being downregulated, likely due to increased protein stability. Supporting this, we observed a cross-species enrichment of proteins involved in protein folding and post-translational processing pathways. In contrast, proteins in Cluster 6 have stable levels despite mRNA being upregulated, suggesting reduced translation or increased proteolysis, which is enriched for *S. enterica* Typhimurium. Furthermore, we observed an enrichment in protein transport and secretion pathway for all species within this cluster. This indicates that the observed protein stability is not only due to translation and degradation activity, but also to a rapid export from the cell. Moreover, genes encoding proteins involved in protein phosphorylation were commonly enriched for the three species in Cluster 4 and 5. In cluster 4, protein levels were upregulated despite stable mRNA abundance, whereas Cluster 5 exhibited mRNAs upregulation and proteins downregulation (**Supplementary Figure 8)**. Notably, Cluster 4 was also commonly enriched for lipid metabolism across the three species. This indicates presence of post-transcriptional regulations such as regulation via small regulatory RNAs involved in carbon metabolism (48, 49) or post-translational modifications which have high prevalence in bacterial metabolic enzymes (50).

### mRNA-protein level stress-specific stimulon genes under stress

We previously identified stress-specific stimulon genes as sets of genes that are uniquely co-regulated in response to distinct stress conditions (2). To assess whether these stimulon genes exhibit distinct mRNA-protein level correlations compared to other genes, we analyzed mRNA and protein differential expression data across all genes and stress-specific stimulons. No significant differences in mRNA-protein level correlations were observed between stimulon genes and other genes across most conditions except in the case of the osmotic stress-specific stimulon (**Figure 6A**). Notably, genes within this stimulon showed a markedly lower mRNA-protein level correlation across all three pathogens studied.

To investigate this further, we used MOBILE framework (34) to identify osmotic stress specific associations between mRNA and protein levels. Through repeated Lasso regression between paired omics layers, MOBILE enables the identification of condition-specific associations without requiring prior knowledge. The transcriptomic and proteomic datasets with all conditions were used as an input for MOBILE, yielding a list of statistically robust associations between proteins and mRNAs. Next, the osmotic stress (Oss) condition data from both datasets were excluded from the input, and MOBILE is re-run to obtain condition-dependent associations list by comparing the all-data run to the latter. Similarly, the control (Ctrl) condition data is held out, and MOBILE re-run yielded another list of associations to identify CTRL-data dependent associations (**Figure 6B**). We identified Oss specific associations by over representation analysis (ORA), comparing the osmotic stress-dependent and control-dependent association lists. Using Oss specific associations gene list, we ran a GO enrichment analysis. Interestingly, translation-related processes and components of the translational machinery were enriched only for *Y. pseudotuberculosis* (**Figure 6C**). This suggests a functional link between osmotic stress-specific mRNA-protein decoupling and translational regulation. In contrast, *S. aureus* showed no significant enrichment in osmotic-specific pathways. This might reflect the inherent physiological resilience of Gram-positive bacteria and metabolic adaptation to osmotic stress (51).

Based on this finding, we hypothesized that the reduced mRNA-protein correlation observed for osmotic stress-specific stimulon genes might result from impaired translation under osmotic stress. To test this, we performed polysome profiling on *Y. pseudotuberculosis* exposed to Oss for 10 minutes, using unexposed bacteria collected at matched time points as controls The polysome profiles showed no notable differences between the osmotic stress and control conditions, with a clear retention of polysome peaks (**Figure 6D**). This indicates that under osmotic stress, ribosomes are still successfully recruited to and load onto transcripts. However, intact polysomes do not necessarily equate to active protein synthesis, as ribosomes may be stalled mid-translation. Therefore, global translational activity may be insufficient to meet the heightened translational demand triggered by the strong transcriptional induction of osmotic stress-specific stimulon genes.

To further characterize the translational response, translation efficiency was assessed using a puromycin incorporation assay across osmotic stress, low iron, and unexposed control conditions. Chloramphenicol-treated bacteria served as a translationally inactive control. Since puromycin causes a premature translation termination, its incorporation serves as a direct proxy for active translation. The results showed that translational activity was significantly reduced under osmotic stress compared to both unexposed control and low iron conditions, indicating that osmotic stress specifically impairs translation activity (**Figure 6E and Supplementary Figure 9**). Therefore, the reduced mRNA-protein correlation for osmotic stress-specific stimulons is not surprising given the overall reduction in translational activity. These stimulon genes exhibit significant changes in mRNA abundance during stress. However, because the translational machinery is impaired, it cannot efficiently process or respond to these altered transcript levels. As a result, the correlation between mRNA and protein decreases. Notably, this decoupling is specific to osmotic stress and was not observed under low iron conditions.

## Discussion

We conducted a comprehensive analysis by integrating transcriptomic and proteomic datasets of *S. enterica* Typhimurium, *Y. pseudotuberculosis*, and *S. aureus* under 10 infection-relevant stress conditions, as well as a control condition during exponential growth. The inclusion of diverse pathogens, which are distantly related yet possess conserved genes, enabled us to identify both common and species-specific mRNA-protein relationships across a range of conditions mimicking host environments. Consistent with prior findings in other organisms, our analyses indicate a positive correlation between mRNA and protein levels across the three bacterial pathogens. However, this correlation is significantly reduced under stress conditions, particularly those that trigger a strong stress response.

This reduction is not surprising, as stress response proteins might be involved in post-transcriptional or post-translational regulations that can decouple mRNA and protein levels. Such regulations can occur through the stabilization or destabilization of mRNA and proteins as a response to the stress. RNA-binding proteins such as CsrA and ProQ can bind to target mRNAs under specific conditions, either stabilizing or destabilizing them, or preventing their translation (52–57). Furthermore, the discoordination of transcription and translation elongation rates under stress conditions may contribute to the decrease in mRNA-protein level correlation (58). For example, in *Escherichia coli* exposed to oxidative stress, there is a significant induction of mRNA expression of the OxyR regulon, which is crucial for survival under oxidative stress, while the translation elongation rate significantly slows down (59, 60).

In contrast, under nutrient-limited conditions, there has been tight coordination between transcription and translation (61, 62). This suggests that the reduced mRNA-protein level correlation in our nutritional downshift condition might not be solely due to the discoordination of transcription and translation but rather other regulatory mechanisms. Conformational changes in riboswitches upon a change in the environment can also lead to reprogrammed mRNA translation and degradation, as is the case for the lysC riboswitch in the presence and absence of lysine (63).

Even though mostly studied in eukaryotes, post-transcriptional nucleotide modifications triggered by stress conditions can also have an impact on mRNA-protein level as it is shown for increased level of m^6^A methylation under hypoxic for human cancer cells (64). On the other hand, m^6^A methylation was shown to be lowered under increased temperature for *P. aeruginosa* (65). Similarly post-translational modifications such as methylation, phosphorylation, acetylation, and sumoylation can directly impact protein stability or indirectly through triggered signaling cascade in both eukaryotes and prokaryotes (66, 67). Even though protein stability was not specifically investigated, it was shown that cellular protein phosphorylation was increased during *Streptomyces* differentiation to form spores, which is induced by stress conditions (68).

To assess whether the mRNA-protein discrepancies observed for DEGs are a common feature across species, we identified conserved DEGs across the three pathogens and compared their expression patterns. These conserved genes exhibited similar non-correlating expression patterns under the same stress conditions, suggesting that post-transcriptional and post-translational modifications affecting these genes may be conserved across species. This highlights the evolutionary significance of these regulatory mechanisms in bacterial adaptation to stress.

Among all conditions investigated for stress stimulons, osmotic stress exhibited the most pronounced mRNA-protein decoupling. This impairment likely stems from a compounding effect on both translation initiation and elongation. Initially, the rapid structural remodeling and cell depolymerization required to manage turgor pressure (69, 70) can physically disrupt membrane integrity and the efficiency of transmembrane transporters. This disruption can lead to deficient amino acid uptake, such as methionine, which would directly impair translation initiation due to the requirement of formylmethionine at the start of bacterial protein (21, 22). Concurrently, our experimental validation points to a secondary bottleneck during active synthesis. The maintenance of intact polysomes alongside significantly reduced puromycin incorporation indicates that ribosomes also stall during elongation. Together, these dual impairments in initiation and elongation prevent the translational machinery from efficiently processing the highly induced stress transcripts, resulting in the observed mRNA-protein decoupling.

Our study underscores the importance of integrating transcriptomic and proteomic data to gain a holistic understanding of bacterial responses to infection-relevant stressors. The observed discrepancies between mRNA and protein levels emphasize the need for caution when inferring protein abundance solely from transcriptomic data. The precise post-transcriptional and post-translational mechanisms contributing to the observed mRNA-protein discrepancies remain to be elucidated, as the currently available datasets do not allow their direct identification, representing an important avenue for future investigation.

In conclusion, our findings provide a comprehensive overview of mRNA-protein relationships under stress conditions in three bacterial pathogens, laying the groundwork for further investigations into the regulatory mechanisms mediating bacterial adaptation and survival in hostile host environments. The multi-species, multi-condition nature of this dataset furthermore represents a valuable resource for training predictive models of protein abundance from mRNA levels, particularly under stress conditions where the mRNA-protein relationship deviates substantially from baseline.

## Materials and Methods

### Bacterial strains and growth conditions

*Salmonella enterica* subsp. enterica serovar Typhimurium SL1344, *Staphylococcus aureus* MSSA476, and *Yersinia pseudotuberculosis* serotype O:3 (strain YPIII) species were selected from PATHOgenex database^1^ to analyze transcriptomic and proteomic data.

*Salmonella enterica* SL1344 and *Staphylococcus aureus* MSSA476 strains used in this study did not carry antibiotic resistance markers and were grown in Luria–Bertani (LB) medium (BD Difco™ LB Agar, Lennox) at 37 °C with shaking. The kanamycin-resistant wt *Y. pseudotuberculosis* (YPIII/pIBX) strain was used in this study. Unless otherwise stated, the wild-type strain was grown in LB medium supplemented with 50 µg/mL kanamycin at 26°C with shaking.

### Stress exposure and sample collection

#### Polysome profiling

The wt *Y. pseudotuberculosis* strain was grown overnight in LB, subcultured to an initial optical density (OD_600_) of 0.05 in 200 mL of LB. When the culture reached an OD_600_ of 0.3, the culture was divided into three 50 mL aliquots, each subjected to one of the following conditions for 10 minutes: (1) unexposed control, (2) osmotic stress (0.5 M NaCl), or (3) low-iron stress (250 µM 2,2′-Bipyridyl; Sigma-Aldrich). Cells were harvested by filtration, immediately flash-frozen in liquid nitrogen, and stored at -80°C until lysis.

Frozen cells were lysed by cryogenic grinding in liquid nitrogen using aluminum oxide. Lysis buffer was added during grinding, and the lysate was allowed to thaw gradually throughout the process. The mixture was transferred into a pre-cooled 1.5 mL reaction tube and immediately placed on ice. Lysates were centrifuged at 18,000 x *g* for 5 minutes at 4°C. The resulting supernatants (∼400 µL) were divided into 50 µL aliquots in new, pre-cooled 1.5 mL tubes and stored at ™80°C. Total RNA concentration was quantified using the Qubit fluorometer. Each aliquot was used only once for downstream polysome profiling.

Linear 15–45% sucrose gradients were prepared using 10% and 50% (w/v) sucrose solutions in 50 mM Tris-HCl (pH 7.5) and supplemented with 100 mM NH_4_Cl, 10 mM MgCl_2_, and 6 mM β-mercaptoethanol. Solutions were sterile filtered and gradients generated using the Gradient Master (BioComp Instruments) following the manufacturer’s preprogrammed settings (SW60Ti rotor, short sucrose program, w/v 1st). Prepared gradients were stored at 4°C overnight prior to use.

For polysome analysis, 50 µg of cell lysate was carefully layered onto each gradient without disturbing the interface. Samples were centrifuged at 40,000 rpm for 2 hours and 30 minutes at 4°C in an Optima L-100K ultracentrifuge (Beckman Coulter) equipped with an SW60Ti rotor. Immediately following centrifugation, the sucrose gradients were carefully placed on ice to prevent disturbance, and continuous polysome absorbance profiles at 254 nm were recorded using the BioComp gradient fractionation system complemented with a TRIAX flow cell. 14 discrete fractions were sequentially collected from the top to the bottom of each gradient using the automated fraction collector, following the manufacturer’s instructions and stored at ™80°C until further analysis.

#### Puromycin assay

To assess active translation, the wt strain was cultured as described above for overnight and subculture preparation. Upon reaching exponential phase (OD_600_ of 0.4), the culture was distributed into four separate flasks representing the following treatment conditions: (1) chloramphenicol (50 µg/mL), (2) unexposed control, (3) osmotic stress (0.5 M NaCl), or (4) low-iron stress (250 µM 2,2′-Bipyridyl; Sigma-Aldrich). Cultures grew with their respective conditions for 10 minutes. Subsequently, to abruptly freeze and capture the translational landscape, both 10 µg/mL puromycin and 200 µg/mL chloramphenicol were simultaneously added to the culture, followed by an additional 10-minute incubation period prior to cell lysis. The experiment was performed in four independent biological replicates.

To extract proteins, 20 mL of bacterial culture was collected from each treatment group and cells were immediately harvested by centrifugation at 4,000 × *g* for 10 minutes at 4°C. The resulting cell pellets were washed twice with phosphate-buffered saline and resuspended in 1 ml of lysis buffer (100 mM Tris-HCl, 10 mM EDTA, 1x cOmplete™ protease inhibitor cocktail (Roche)). The cell suspensions were lysed on ice using sonication for nine 20 second cycles, each cycle comprised

20 second sonication step separated by 30 second intervals with 10% amplitude. Following centrifugation at 13,000 x *g* for 10 minutes at 4°C, the supernatant was collected and stored at - 20°C. Protein concentrations were defined using QuBit Protein Assay Kit (Catalog Number: Q33211).

#### Immunoblotting

Protein concentrations were fixed across samples, and proteins were mixed in 1x sample buffer (2% SDS, 62.5 mM Tris-HCl, 20% glycerol, 0.05% bromophenol blue, 1% β-mercaptoethanol). 17 µl of proteins were separated by sodium dodecyl sulfate (SDS)-PAGE on 4-15% Mini-PROTEAN TGX gels using 80V for 20 minutes, followed by 120V for 1 hour. SDS gel was transferred onto methanol activated Immobilon-P PVDF membrane (Millipore) by semidry electroblotting using Bio-Rad Trans-Blot SD Semi-Dry Transfer Cell (70V for 1 hour 10 minutes). Membranes were blocked with 5% milk in 1x Tris-buffered saline with Tween (TBST) and probed with Anti-Puromycin at 1:10,000 dilution (Sigma-Aldrich, MABE343, clone 12D10) primary antibody in 1x TBST, overnight at 4°C. Following each antibody incubation, the membranes were washed three times for 10 minutes with 1x TBST. Membranes were incubated with secondary antibody: monoclonal Sheep Anti-Mouse IgG - Horseradish Peroxidase at 1:7,000 dilution (Cytiva, NA931V) in 1x TBST for 1 hour at room temperature. The final blot was developed by chemiluminescence (Immobilon Western Chemiluminescent HRP Substrate, Millipore) for 5 minutes, and the bands were detected with Amersham Imager 680. Band intensities were quantified using ImageJ (version 1.54p).

#### Transcriptomic data retrieval and proteomic data generation

The transcriptome data published for *S. enterica, Y. pseudotuberculosis*, and *S. aureus* were obtained from Gene Expression Omnibus (GEO) (71) with accession number GSE152295.

Proteomics data was obtained by exposing the three bacterial species to the same stress conditions as mentioned in Avican *et al*. (1), with unexposed control condition and excluding virulence inducing condition. After stress exposures, 3 ml of bacterial cultures were pelleted with 9000 RPM for 2 minutes centrifugation and cell pellets were lysed in 200 µl 2% SDS. Thereafter, 5 μg of each sample were reduced with 20 mM TCEP. Samples were digested with a modified sp3 protocol (72), as previously described (42). Briefly, samples were added to a bead suspension (10 μg of beads (Sera-Mag Speed Beads, 4515-2105-050250, 6515-2105-050250) in 10 μl 15% formic acid and 30 μl ethanol) and incubated shaking for 15 min at room temperature. Beads were then washed four times with 70% ethanol. Proteins were digested overnight by adding 40 μl of 5 mM chloroacetamide, 1.25 mM TCEP, and 200 ng trypsin in 100 mM HEPES pH 8.5. Peptides were eluted from the beads and were dried under vacuum. Peptides were then labelled with TMT11plex (Thermo Fisher Scientific), pooled and desalted with solid-phase extraction using a Waters OASIS HLB μElution Plate (30 μm). Samples were fractionated into 48 fractions on a reversed-phase C18 system running under high pH conditions, with every sixth fraction being pooled together. Samples were analyzed by LC-MS/MS using a data-dependent acquisition strategy on a Thermo Fisher Scientific Vanquish Neo LC coupled with an Thermo Fisher Scientific Orbitrap Exploris 480. Raw files were processed with MSFragger using standard settings for TMT (73) against FASTA databases of each organism obtained from NCBI; for *S. enterica* Typhimurium SL1344 accession numbers NC_016810, NC_017718, NC_017719, NC_017720; for *Y. pseudotuberculosis* accession number NC_010465; for *S. aureus* (MSSA476) accession number NC_002952. The mass spectrometry proteomics data have been deposited to the ProteomeXchange Consortium via the PRIDE partner repository (74) with the dataset identifier PXD055243.

### Data processing and normalization

#### Processing of transcriptomic data

Transcriptomic data were imported from PATHOgenex, including statistical values such as LogFC, *p*-value, and adjusted *p*-value. Moreover, the differential expression analysis was also already conducted, and thus no further pre-processing was done here.

#### Processing of proteomic data

Raw protein files containing intensity data from TMT runs were imported into R (75) (version 4.4.2). Data were pre-processed to retain only protein locus tag, intensities values, and razor peptide count -defined as peptides shared across multiple proteins. Proteins were then filtered to retain only those quantified by at least two unique peptides and exhibiting non-zero intensity values; entries with zero intensity were considered missing data and excluded. This processing yielded one matrix per species per run, in which each row contained a protein identifier, its total intensity, and its corresponding razor peptide count.

To estimate protein abundance, each protein was mapped to its corresponding treatment condition. The relative reporter ion abundance of a given protein for its treatment was divided by the total relative reporter ion abundance as a first estimate of protein abundance. Given the large variation in protein length observed across the dataset — ranging from 10 to 3,000 amino acids in *S. enterica* and *Y. pseudotuberculosis*, and from 10 to approximately 2,000 amino acids in *S. aureus* (**Supplementary Figure 10**), we applied a correction to account for size-dependent differences in peptide yield. Since longer proteins produce more tryptic peptides upon digestion, and consequently more ions, the initial estimate was divided by the number of expected tryptic peptides to obtain iBAQ (15). This ensures that iBAQ reflects protein abundance independently of protein size. Matrices from the same species were subsequently merged across runs, and proteins absent across all replicates were excluded, yielding three species-specific matrices each containing iBAQ values for every detected protein across the three biological replicates and their associated treatment conditions.

To identify the most appropriate normalization strategy, 8 normalization methods were evaluated using the NormalyzerDE (44) package (version 1.23.2): VSN, Log2-transformation, Quantile, Global Intensity, Mean, Median, Linear Regression, and Cyclic Loess. Normalized outputs were visualized using principal component analysis (PCA) plots generated with ggplot2 (76) (version 4.0.2) to assess replicate clustering within each stress condition. VSN consistently produced the tightest replicate clustering across condition and was therefore applied to the three species-specific matrices to generate VSN-normalized iBAQ values for downstream analysis.

Differential abundance analysis was performed on the normalized proteome data using the limma (45) package (version 3.58.1) with default parameters, applying empirical Bayes moderation to stabilize variance estimates across proteins. Differentially abundant proteins were identified for each species to characterize the proteomic response to every stress condition, and highlight proteins potentially involved in key regulatory mechanisms.

## Data analysis

### Correlation analysis

To assess the correlation between mRNA and protein levels, we used the log2-transformed mean TPM values for mRNAs and the log2-transformed mean iBAQ values for proteins from three replicates of each condition. The Pearson correlation coefficient was calculated for each condition to determine the strength of the correlation. Statistical significance p-value < 0.05 was evaluated using Student’s t-test.

### Identification of gene clusters with discordant mRNA and protein level regulation

To identify genes with discordant mRNA and protein level regulation under 10 stress conditions, the Z-score values of mRNA from control data were combined with the Z-score values of protein, calculated using limma differential expression analysis to create a single data table and visualize using scatter plots separately for each species. Thereafter, the combined transcriptome and proteome data were filtered with *p*-value < 0.05 to select genes whose mRNA or protein level changes upon exposure to stress conditions.

Genes were then grouped in 6 different clusters based on the expression patterns as described in Table 1. Cluster 1; mRNA downregulated in at least one stress condition but protein level is not changed, Cluster 2; mRNA downregulated in at least one condition but protein is upregulated, Cluster 3; mRNA level is not changed in any condition but protein is downregulated, Cluster 4; mRNA level is not changed in any condition but protein is upregulated in at least one condition, Cluster 5; mRNA is upregulated in at least one condition but protein is downregulated, and Cluster 6; mRNA is upregulated in at least one condition but protein level is not changed.

### Integrative omics analysis with MOBILE

MOBILE pipeline (34) is used for three species (SA, SE, and YP) with 11 samples including both transcriptomic and proteomic datasets. The iBAQ datasets were log2-transformed and normalized. First, they were filtered to only retain the most highly variant features (top 10% for transcripts and 20% for proteins). Next, the MOBILE pipeline (MATLAB version R2023a) is run, using the glmnet package (77) and *Y* = *β. X* + *δ* formalism, where *Y* matrix is the proteomics dataset and *X* is the transcriptomics. The objective is to obtain the coefficients matrix *β. δ* is the y-intercept values, which are almost zero by design. The regression process is repeated 10000 times with randomly selected random number generation seeds to yield an ensemble of Lasso matrices. By identifying coefficients that appeared in at least half of the models, a robust Lasso coefficient matrix is selected. This matrix represents the stable associations between analytes across omics layers.

To find stress-specific interactions, MOBILE employs a “Leave-One-Group-Out” (LOGO) strategy where each sample exposed to the stress is excluded from the input one-at-a-time. Here, both the osmotic stress and control condition samples were subtracted, and corresponding final matrices were named as OSS-IAN and CTRL-IAN matrices. IANs are the Integrated Association Networks that are coalesced gene-level networks, where nodes represent genes of the matrix analytes and edges stand for the non-zero coefficients.

### Functional annotation

#### Enrichment analysis

Gene Ontology (GO) enrichment analyses were conducted using the clusterProfiler (78) package v4.10.1 in R. Most significant enriched terms were identified using a cut-off criterion of p-value<0.05. Plots showing the most significant enriched Biological Process (BP) GO terms were generated using the ggplot2 package.

#### Over representation analysis (ORA)

The subsequent step involves comparing OSS-IAN and FULL-IAN (all condition matrix), to obtain OSS-dependent interactions. Similarly, comparing CTRL-IAN to FULL-IAN yields CTRL-dependent interactions. Next, pathway enrichment analysis (clusterProfiler version 4.14.6) was adopted for the genes list obtained by comparing OSS-dependent list to CTRL-dependent gene list. Gene Ontology gene sets curated for three species (*SA, SE, YP)* were used in gmt format with a minimum gene set size of 15 and a maximum of 500. Significantly enriched gene sets were filtered by q-value < 0.2 and p-value < 0.05.

## Data availability

All data necessary to reproduce the findings of this study are publicly available without restriction.

The complete dataset is available from Zenodo (79) (https://zenodo.org/records/19488690). Transcriptomic data were obtained from PATHOgenex RNA Atlas (1); Protein sequences and Mass spectrometry results are provided in the Zenodo repository.

## Code Availability

All code and commands to reproduce the analysis are included in a publicly available.

A repository containing a Rnotebook with all the code to reproduce the figures and analyzes presented in this manuscript, is available on GitHub at https://github.com/avicanlab/RNAProtCorr.

All packages and versions are listed in the Renv file available in the GitHub repository.

## References

1. K. Avican, et al., RNA atlas of human bacterial pathogens uncovers stress dynamics linked to infection. Nat Commun 12, 3282 (2021).

2. L. Fernandez, M. Rosvall, J. Normark, M. Fallman, K. Avican, Co-PATHOgenex web application for assessing complex stress responses in pathogenic bacteria. Microbiol Spectr 12, e02781–23 (2024).

3. M. J. Lee, M. B. Yaffe, Protein Regulation in Signal Transduction. Cold Spring Harb Perspect Biol 8, a005918 (2016).

4. J. Bathke, A. Konzer, B. Remes, M. McIntosh, G. Klug, Comparative analyses of the variation of the transcriptome and proteome of Rhodobacter sphaeroides throughout growth. BMC Genomics 20, 358 (2019).

5. Y. Zhu, et al., Integrative analysis of transcriptome and proteome provides insights into adaptation to cadmium stress in Sedum plumbizincicola. Ecotoxicol Environ Saf 230, 113–149 (2022).

6. S. Liu, et al., Transcriptome and Proteome of Methicillin-Resistant Staphylococcus aureus Small-Colony Variants Reveal Changed Metabolism and Increased Immune Evasion. Microbiol Spectr 11, e0189822 (2023).

7. F. Grunberger, et al., Uncovering the temporal dynamics and regulatory networks of thermal stress response in a hyperthermophile using transcriptomics and proteomics. mBio 14, e0217423 (2023).

8. F. Edfors, et al., Gene-specific correlation of RNA and protein levels in human cells and tissues. Mol Syst Biol 12, 883 (2016).

9. X. Feng, Q. Ma, Transcriptome and proteome profiling revealed molecular mechanism of selenium responses in bread wheat (Triticum aestivum L.). BMC Plant Biol 21, 584 (2021).

10. M. Backman, et al., Multi-omics insights into functional alterations of the liver in insulin-deficient diabetes mellitus. Mol Metab 26, 30–44 (2019).

11. G. Chen, et al., Discordant protein and mRNA expression in lung adenocarcinomas. Mol Cell Proteomics 1, 304–13 (2002).

12. C. Vogel, E. M. Marcotte, Insights into the regulation of protein abundance from proteomic and transcriptomic analyses. Nat Rev Genet 13, 227–32 (2012).

13. F. Gebauer, M. W. Hentze, Molecular mechanisms of translational control. Nat Rev Mol Cell Biol 5, 827–835 (2004).

14. T. Maier, M. Güell, L. Serrano, Correlation of mRNA and protein in complex biological samples. FEBS Letters 583, 3966–3973 (2009).

15. B. Schwanhäusser, et al., Global quantification of mammalian gene expression control. Nature 473, 337–342 (2011).

16. M. Jovanovic, et al., Dynamic profiling of the protein life cycle in response to pathogens. Science 347, 1259038 (2015).

17. M. Zhang, et al., Impact of Growth Rate on the Protein-mRNA Ratio in Pseudomonas aeruginosa. mBio 14, e0306722 (2023).

18. M. U. Caglar, et al., The E. coli molecular phenotype under different growth conditions. Sci Rep 7, 45303 (2017).

19. Y. W. Choi, S. A. Park, H. W. Lee, D. S. Kim, N. G. Lee, Analysis of growth phase-dependent proteome profiles reveals differential regulation of mRNA and protein in Helicobacter pylori. PROTEOMICS 8, 2665–2675 (2008).

20. W.-H. Chen, et al., Integration of multi-omics data of a genome-reduced bacterium: Prevalence of post-transcriptional regulation and its correlation with protein abundances. Nucleic Acids Res 44, 1192–1202 (2016).

21. K. Becker, et al., Quantifying post-transcriptional regulation in the development of Drosophila melanogaster. Nat Commun 9, 4970 (2018).

22. R. W. Corbin, et al., Toward a Protein Profile of Escherichia coli: Comparison to Its Transcription Profile. Proceedings of the National Academy of Sciences of the United States of America 100, 9232–9237 (2003).

23. S. P. Gygi, Y. Rochon, B. R. Franza, R. Aebersold, Correlation between Protein and mRNA Abundance in Yeast. Molecular and Cellular Biology 19, 1720–1730 (1999).

24. L. Jeacock, J. Faria, D. Horn, Codon usage bias controls mRNA and protein abundance in trypanosomatids. eLife 7, e32496 (2018).

25. L. Ponnala, Y. Wang, Q. Sun, K. J. van Wijk, Correlation of mRNA and protein abundance in the developing maize leaf. The Plant Journal 78, 424–440 (2014).

26. J. Wang, et al., Combined proteomic and transcriptomic analysis of the antimicrobial mechanism of tannic acid against Staphylococcus aureus. Front Pharmacol 14, 1178177 (2023).

27. S. Kavela, et al., Use of an Integrated Multi-Omics Approach To Identify Molecular Mechanisms and Critical Factors Involved in the Pathogenesis of Leptospira. Microbiol Spectr 11, e0313522 (2023).

28. L. Huang, et al., Integration of Transcriptomic and Proteomic Approaches Reveals the Temperature-Dependent Virulence of Pseudomonas plecoglossicida. Front Cell Infect Microbiol 8, 207 (2018).

29. C. Kocharunchitt, T. King, K. Gobius, J. P. Bowman, T. Ross, Integrated transcriptomic and proteomic analysis of the physiological response of Escherichia coli O157:H7 Sakai to steady-state conditions of cold and water activity stress. Mol Cell Proteomics 11, M111.009019 (2012).

30. Q. Xiong, et al., Integrated transcriptomic and proteomic analysis of the global response of Synechococcus to high light stress. Mol Cell Proteomics 14, 1038–53 (2015).

31. F. J. Perez-Llarena, G. Bou, Proteomics As a Tool for Studying Bacterial Virulence and Antimicrobial Resistance. Front Microbiol 7, 410 (2016).

32. X. Yang, et al., RNA markers for ultra-rapid molecular antimicrobial susceptibility testing in fluoroquinolone-treated Klebsiella pneumoniae. J Antimicrob Chemother 75, 1747–1755 (2020).

33. T. Maier, et al., Quantification of mRNA and protein and integration with protein turnover in a bacterium. Mol Syst Biol 7, MSB201138 (2011).

34. C. Erdem, S. M. Gross, L. M. Heiser, M. R. Birtwistle, MOBILE pipeline enables identification of context-specific networks and regulatory mechanisms. Nat Commun 14, 3991 (2023).

35. T. Kwon, H. K. Huse, C. Vogel, M. Whiteley, E. M. Marcotte, Protein-to-mRNA Ratios Are Conserved between Pseudomonas aeruginosa Strains. J. Proteome Res. 13, 2370–2380 (2014).

36. K. P. Jayapal, et al., Uncovering Genes with Divergent mRNA-Protein Dynamics in Streptomyces coelicolor. PLOS ONE 3, e2097 (2008).

37. B. Li, C. N. Dewey, RSEM: accurate transcript quantification from RNA-Seq data with or without a reference genome. BMC Bioinformatics 12, 323 (2011).

38. P. Lu, C. Vogel, R. Wang, X. Yao, E. M. Marcotte, Absolute protein expression profiling estimates the relative contributions of transcriptional and translational regulation. Nat Biotechnol 25, 117–124 (2007).

39. L. Barquist, et al., A comparison of dense transposon insertion libraries in the Salmonella serovars Typhi and Typhimurium. Nucleic Acids Res 41, 4549–4564 (2013).

40. R. R. Chaudhuri, et al., Comprehensive identification of essential Staphylococcus aureus genes using Transposon-Mediated Differential Hybridisation (TMDH). BMC Genomics 10, 291 (2009).

41. S. J. Willcocks, R. A. Stabler, H. S. Atkins, P. F. Oyston, B. W. Wren, High-throughput analysis of Yersinia pseudotuberculosis gene essentiality in optimised in vitro conditions, and implications for the speciation of Yersinia pestis. BMC Microbiol 18, 46 (2018).

42. A. Mateus, et al., The functional proteome landscape of Escherichia coli. Nature 588, 473–478 (2020).

43. O. M. Sigalova, A. Shaeiri, M. Forneris, E. E. Furlong, J. B. Zaugg, Predictive features of gene expression variation reveal mechanistic link with differential expression. Mol Syst Biol 16, MSB209539 (2020).

44. J. Willforss, A. Chawade, F. Levander, NormalyzerDE: Online Tool for Improved Normalization of Omics Expression Data and High-Sensitivity Differential Expression Analysis. J Proteome Res 18, 732–740 (2019).

45. M. E. Ritchie, et al., limma powers differential expression analyses for RNA-sequencing and microarray studies. Nucleic Acids Res 43, e47 (2015).

46. M. S. Robles, J. Cox, M. Mann, In-Vivo Quantitative Proteomics Reveals a Key Contribution of Post-Transcriptional Mechanisms to the Circadian Regulation of Liver Metabolism. PLOS Genetics 10, e1004047 (2014).

47. M. Ashburner, et al., Gene Ontology: tool for the unification of biology. Nat Genet 25, 25–29 (2000).

48. S. Durica-Mitic, Y. Göpel, B. Görke, Carbohydrate Utilization in Bacteria: Making the Most Out of Sugars with the Help of Small Regulatory RNAs. Microbiology Spectrum 6, 10.1128/microbiolspec.rwr-0013–2017 (2018).

49. M. Raina, et al., Dual-function AzuCR RNA modulates carbon metabolism. Proceedings of the National Academy of Sciences 119, e2117930119 (2022).

50. T. Pisithkul, N. M. Patel, D. Amador-Noguez, Post-translational modifications as key regulators of bacterial metabolic fluxes. Current Opinion in Microbiology 24, 29–37 (2015).

51. M. Hecker, J. Pané-Farré, V. Uwe, SigB-Dependent General Stress Response in Bacillus subtilis and Related Gram-Positive Bacteria. Annual Review of Microbiology 61, 215–236 (2007).

52. H. Yakhnin, et al., CsrA Represses Translation of sdiA, Which Encodes the N-Acylhomoserine-l-Lactone Receptor of Escherichia coli, by Binding Exclusively within the Coding Region of sdiA mRNA. Journal of Bacteriology 193, 6162–6170 (2011).

53. C. Pourciau, Y.-J. Lai, M. Gorelik, P. Babitzke, T. Romeo, Diverse Mechanisms and Circuitry for Global Regulation by the RNA-Binding Protein CsrA. Front. Microbiol. 11 (2020).

54. X. Wang, et al., CsrA post-transcriptionally represses pgaABCD, responsible for synthesis of a biofilm polysaccharide adhesin of Escherichia coli. Molecular Microbiology 56, 1648–1663 (2005).

55. A. K. Dubey, et al., CsrA Regulates Translation of the Escherichia coli Carbon Starvation Gene, cstA, by Blocking Ribosome Access to the cstA Transcript. Journal of Bacteriology 185, 4450–4460 (2003).

56. E. Holmqvist, L. Li, T. Bischler, L. Barquist, J. Vogel, Global Maps of ProQ Binding In Vivo Reveal Target Recognition via RNA Structure and Stability Control at mRNA 3′ Ends. Molecular Cell 70, 971–982.e6 (2018).

57. A. Smirnov, C. Wang, L. L. Drewry, J. Vogel, Molecular mechanism of mRNA repression in trans by a ProQ-dependent small RNA. EMBO J 36, 1029–1045 (2017).

58. M. Irastortza-Olaziregi, O. Amster-Choder, Coupled Transcription-Translation in Prokaryotes: An Old Couple With New Surprises. Front. Microbiol. 11 (2021).

59. M. Zhu, X. Dai, Maintenance of translational elongation rate underlies the survival of Escherichia coli during oxidative stress. Nucleic Acids Res 47, 7592–7604 (2019).

60. M. Zhu, X. Dai, Bacterial stress defense: the crucial role of ribosome speed. Cell. Mol. Life Sci. 77, 853–858 (2020).

61. M. Zhu, M. Mori, T. Hwa, X. Dai, Disruption of transcription–translation coordination in Escherichia coli leads to premature transcriptional termination. Nat Microbiol 4, 2347–2356 (2019).

62. U. Vogel, K. F. Jensen, The RNA chain elongation rate in Escherichia coli depends on the growth rate. Journal of Bacteriology 176, 2807–2813 (1994).

63. M.-P. Caron, et al., Dual-acting riboswitch control of translation initiation and mRNA decay. Proceedings of the National Academy of Sciences 109, E3444–E3453 (2012).

64. B. W. Scheithauer, J. M. Bruner, Central nervous system tumors. Clin Lab Med 7, 157–179 (1987).

65. X. Deng, et al., Widespread occurrence of N6-methyladenosine in bacterial mRNA. Nucleic Acids Res 43, 6557–6567 (2015).

66. J. M. Lee, H. M. Hammarén, M. M. Savitski, S. H. Baek, Control of protein stability by post-translational modifications. Nat Commun 14, 201 (2023).

67. J. A. Cain, N. Solis, S. J. Cordwell, Beyond gene expression: The impact of protein post-translational modifications in bacteria. Journal of Proteomics 97, 265–286 (2014).

68. A. Manteca, J. Ye, J. Sánchez, O. N. Jensen, Phosphoproteome Analysis of Streptomyces Development Reveals Extensive Protein Phosphorylation Accompanying Bacterial Differentiation. J. Proteome Res. 10, 5481–5492 (2011).

69. E. Bremer, R. Krämer, Responses of Microorganisms to Osmotic Stress. Annu. Rev. Microbiol. 73, 313–334 (2019).

70. E. A. Mueller, P. A. Levin, Bacterial Cell Wall Quality Control during Environmental Stress. mBio 11, 10.1128/mbio.02456-20 (2020).

71. T. Barrett, et al., NCBI GEO: archive for functional genomics data sets—update. Nucleic Acids Res 41, D991–D995 (2013).

72. C. S. Hughes, et al., Ultrasensitive proteome analysis using paramagnetic bead technology. Mol Syst Biol 10, 757 (2014).

73. A. T. Kong, F. V. Leprevost, D. M. Avtonomov, D. Mellacheruvu, A. I. Nesvizhskii, MSFragger: ultrafast and comprehensive peptide identification in mass spectrometry-based proteomics. Nat Methods 14, 513–520 (2017).

74. E. W. Deutsch, et al., The ProteomeXchange consortium at 10 years: 2023 update. Nucleic Acids Res 51, D1539–D1548 (2023).

75. R Core Team, R: A Language and Environment for Statistical Computing (R Foundation for Statistical Computing, 2024).

76. H. Wickham, ggplot2: Elegant Graphics for Data Analysis (Springer-Verlag New York, 2016).

77. J. H. Friedman, T. Hastie, R. Tibshirani, Regularization Paths for Generalized Linear Models via Coordinate Descent. Journal of Statistical Software 33, 1–22 (2010).

78. T. Wu, et al., clusterProfiler 4.0: A universal enrichment tool for interpreting omics data. Innovation (Camb) 2, 100141 (2021).

79. European Organization For Nuclear Research, OpenAIRE, Zenodo: Research. Shared. [Preprint] (2013). Available at: https://www.zenodo.org/ x[Accessed 11 December 2025].

