## Supplementary Information for "Bacterial Stress Responses Lower mRNA-Protein Level Correlations"

X first author

*Correspondence: Kemal Avican

**This PDF file includes:**

Supporting text
Figures S1 to S10
Tables S1 to S3

### Figures

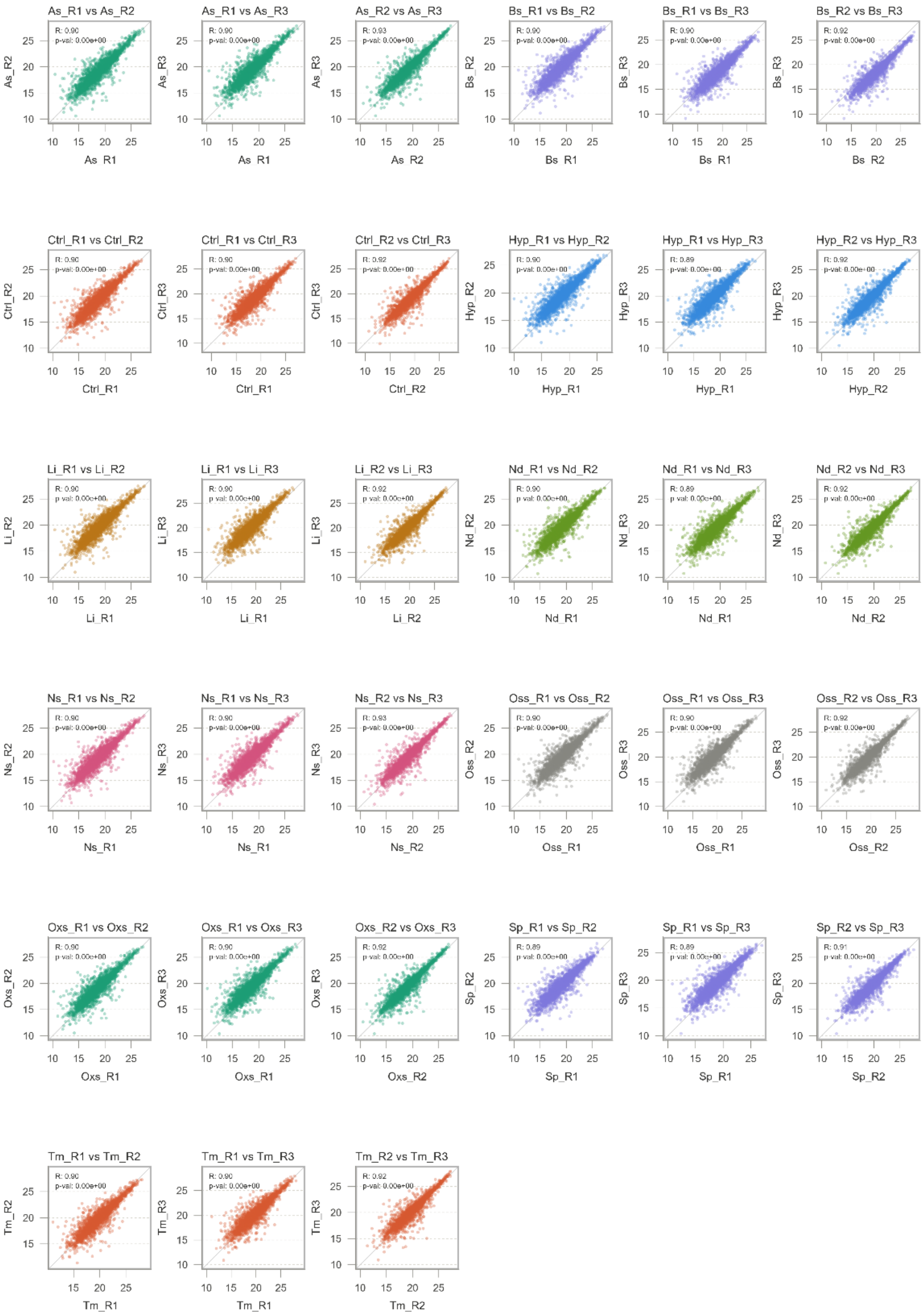


**Fig. S1. Replicates each condition show similar global expression patterns in *S. enterica* Typhimurium.** Pairwise comparison of $log_2$-transformed protein expression values (iBAQ) from each of the three replicates per condition were performed using proteomic analysis. The Pearson correlation coefficient (R) was used to measure similarity, and the Student's T-test was used to calculate the *p*-value.

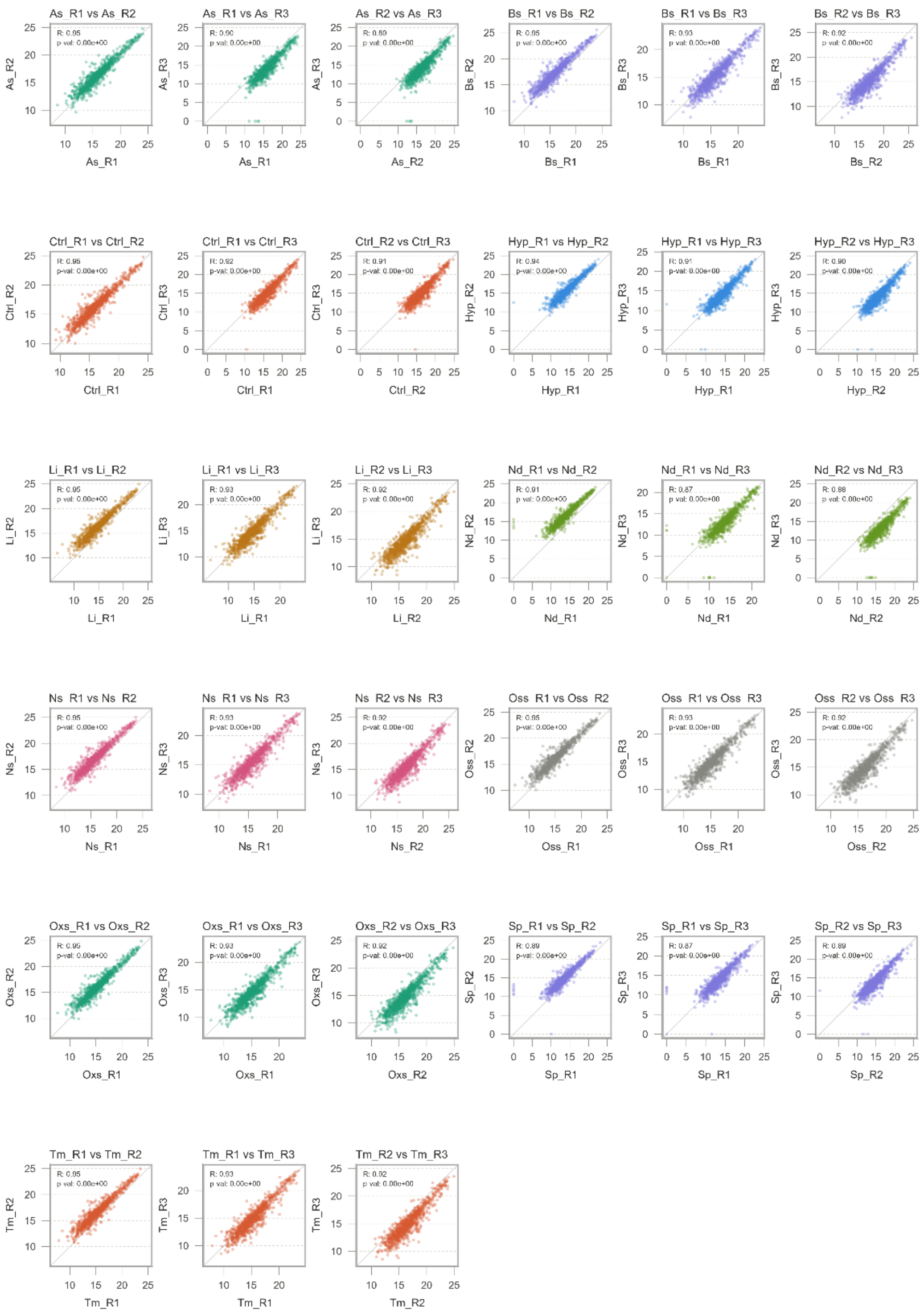


**Fig. S2. Replicates each condition show similar global expression patterns in *S. aureus*.** Pairwise comparison of $\log_2$-transformed protein expression values (iBAQ) from each of the three replicates per condition were performed using proteomic analysis. The Pearson correlation coefficient (R) was used to measure similarity, and the Student's T-test was used to calculate the *p*-value.

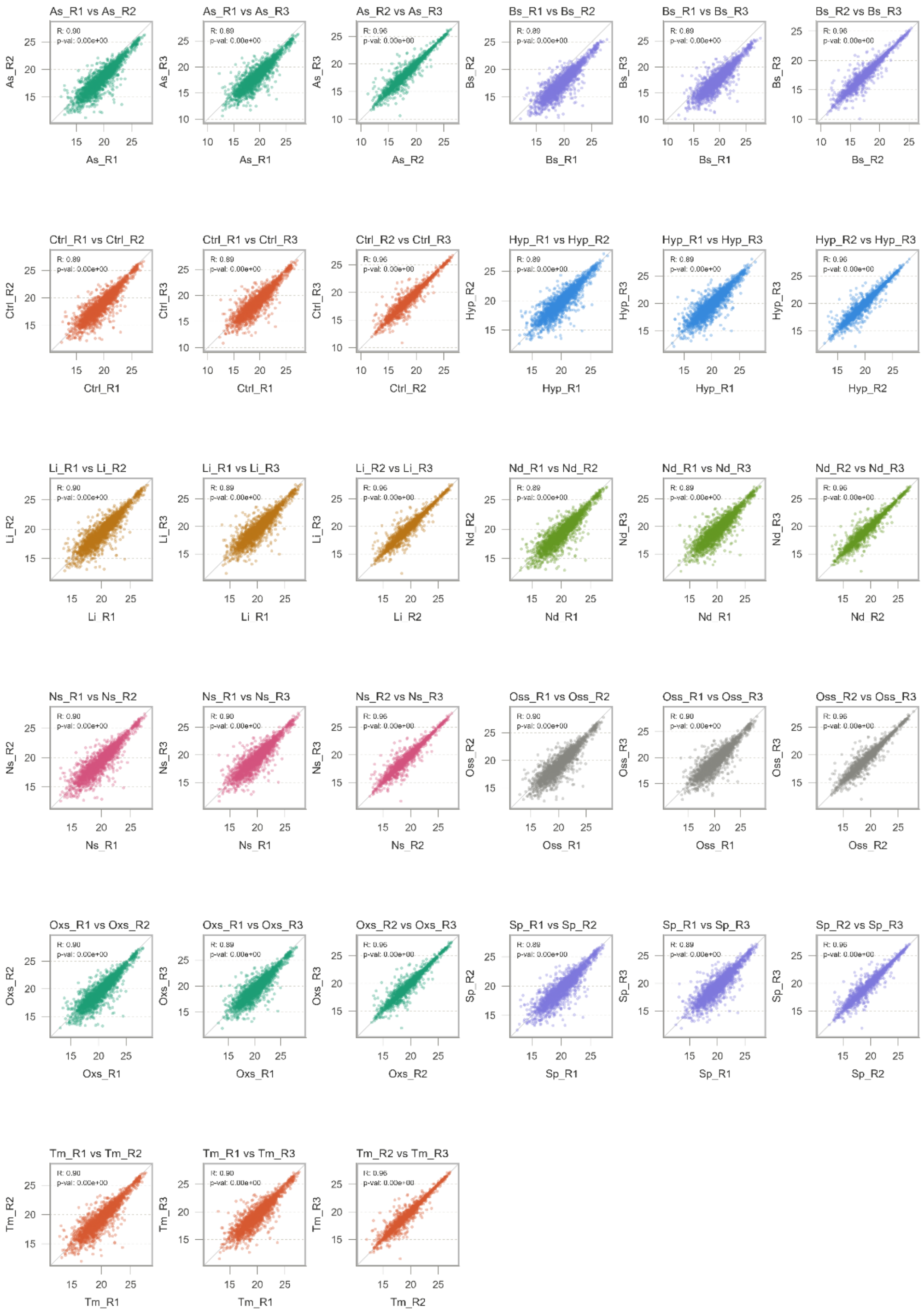

**Fig. S3. Replicates each condition show similar global expression patterns in *Y. pseudotuberculosis*.** Pairwise comparison of $log_2$-transformed protein expression values (iBAQ) from each of the three replicates per condition were performed using proteomic analysis. The Pearson correlation coefficient (R) was used to measure similarity, and the Student's T-test was used to calculate the *p*-value.

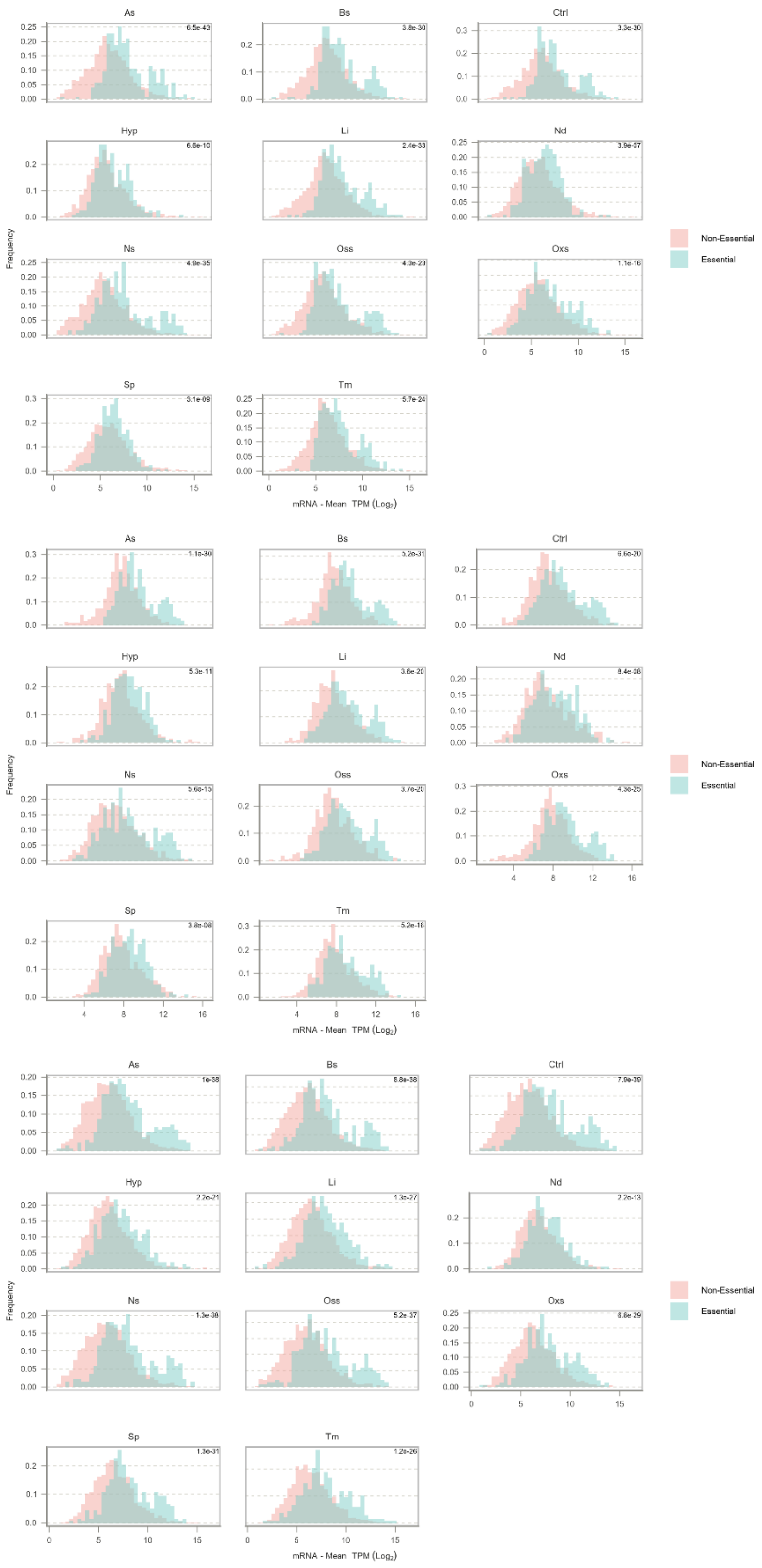


**Fig. S4. mRNA expression of essential genes are higher than non-essential genes in all tested conditions for the three species. A)** Frequency of gene expression levels in $log_2$-transformed TPM values in *S. enterica* Typhimurium **B)** *S. aureus* **C)** *Y. pseudotuberculosis*.

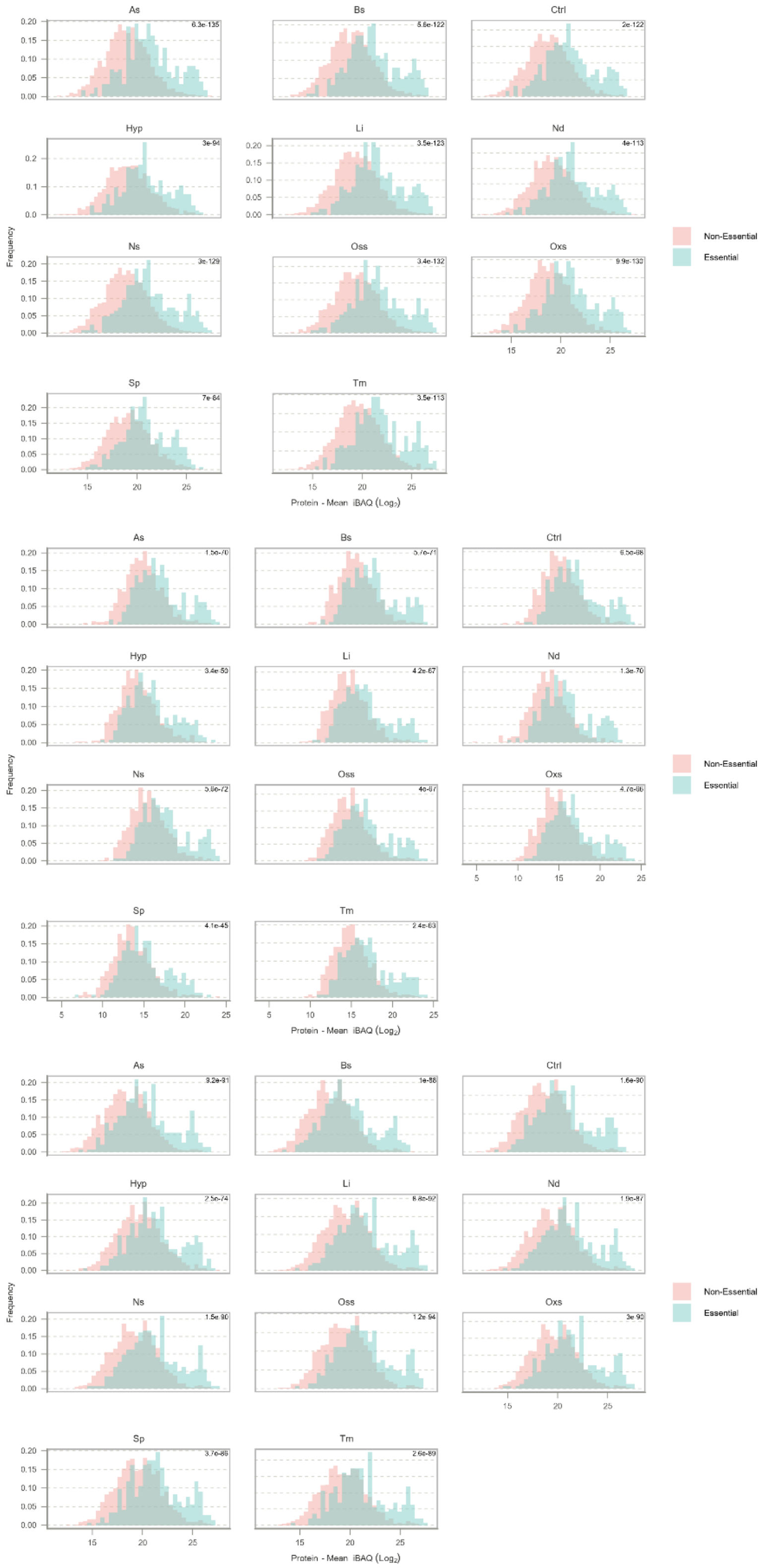


**Fig. S5. Protein expression of essential genes are higher than non-essential genes in all tested conditions for the three species. A)** Frequency of gene expression levels in $log_2$-transformed iBAQ values in *S. enterica* Typhimurium **B)** *S. aureus* **C)** *Y. pseudotuberculosis*.

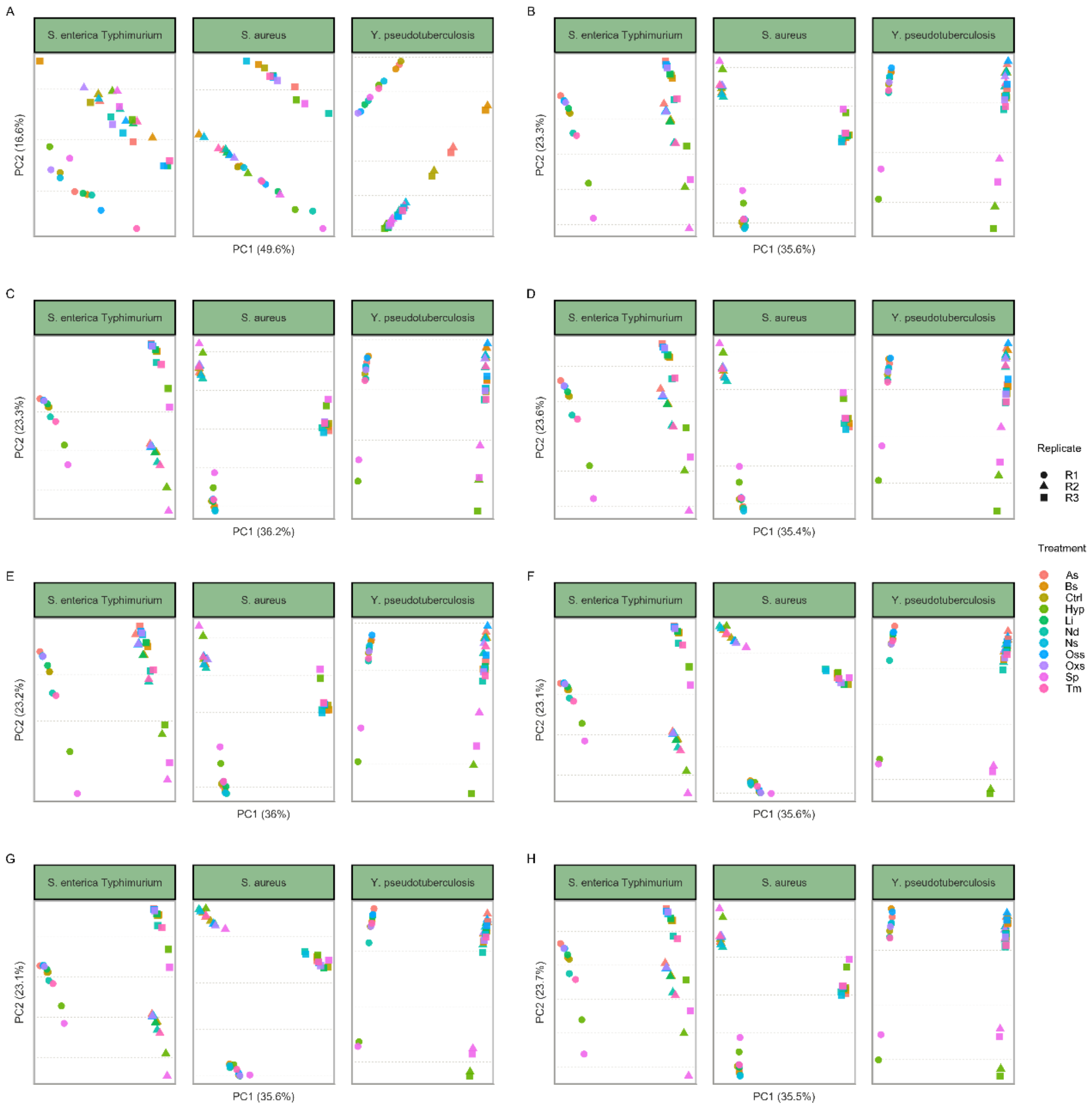


**Fig. S6. Variance Stabilizing Normalization (VSN) method performs better in clustering replicates from the same condition and separation between different conditions for proteome data from *S enterica* Typhimurium, *S. aureus*, *Y. pseudotuberculosis*.** Principle Component Analysis (PCA) results of proteome data using different normalization methods are shown in panels for: **A)** Log2, **B)** VSN, **C)** CycLoess, **D)** Quantile, **E)** Median, **F)** Mean, **G)** Global Intensity, **H)** Linear regression.

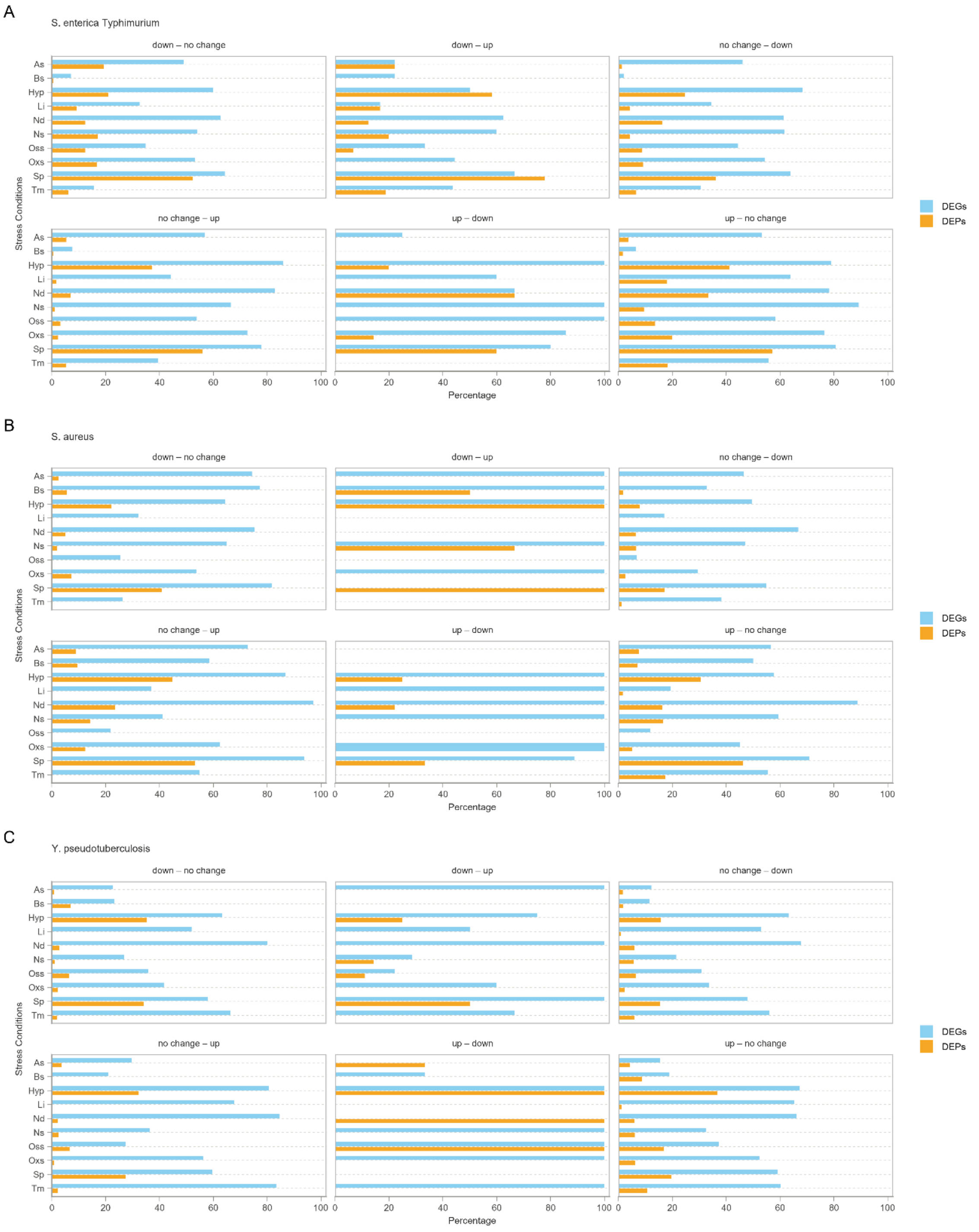


**Fig. S7. Genes with discordantly regulated mRNA and protein levels are found in all tested stress conditions for the three species.** Percentage of DEGs and DEPs, filtered with *p*-values<0.05, in each of the 6 discordantly regulated gene clusters for every tested stress conditions are shown as bars for **A)** *S. enterica* Typhimurium, **B)** *S. aureus*, **C)** *Y. pseudotuberculosis.*

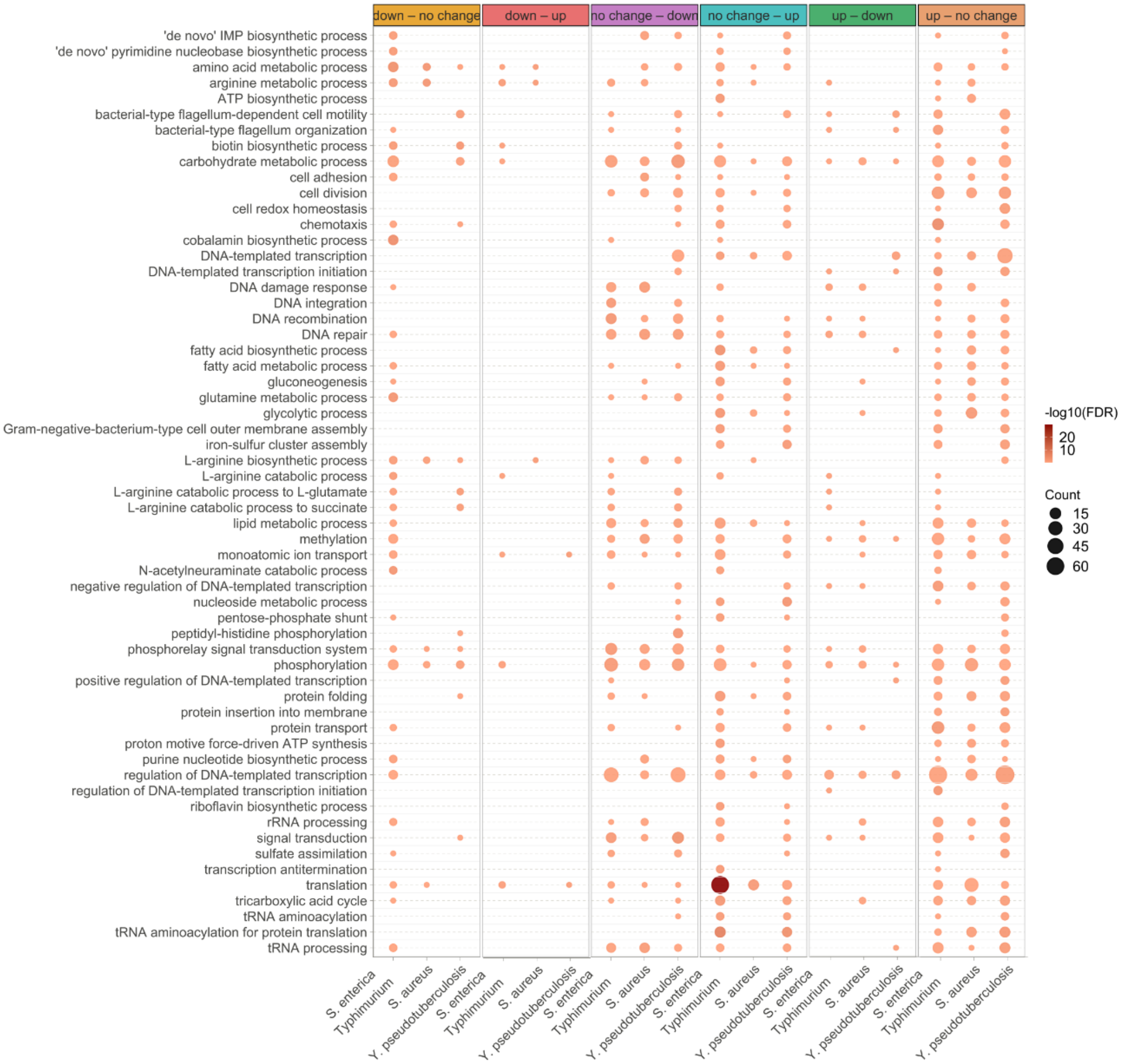


**Fig. S8 Stress response induces different patterns of discordant mRNA and protein level regulation highlighting key role for translation.** GO biological process enrichment analysis of 6 discordantly regulated gene clusters for the three species.

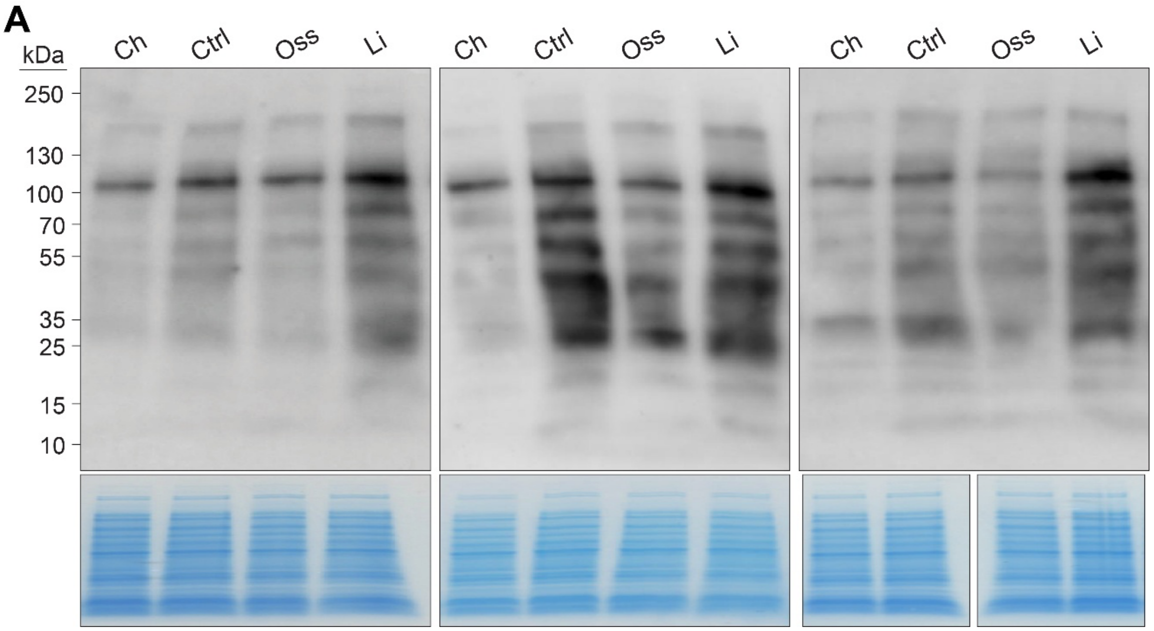


**Fig. S9. Translation efficiency is reduced under osmotic stress across biological replicates. A)** Western blot membranes showing active translation of nascent proteins detected using monoclonal anti-puromycin antibody in wt *Y. pseudotuberculosis* cells under negative control (Ch), positive control (Ctrl), osmotic stress (Oss), and low iron (Li) conditions. Three independent biological replicates are shown.

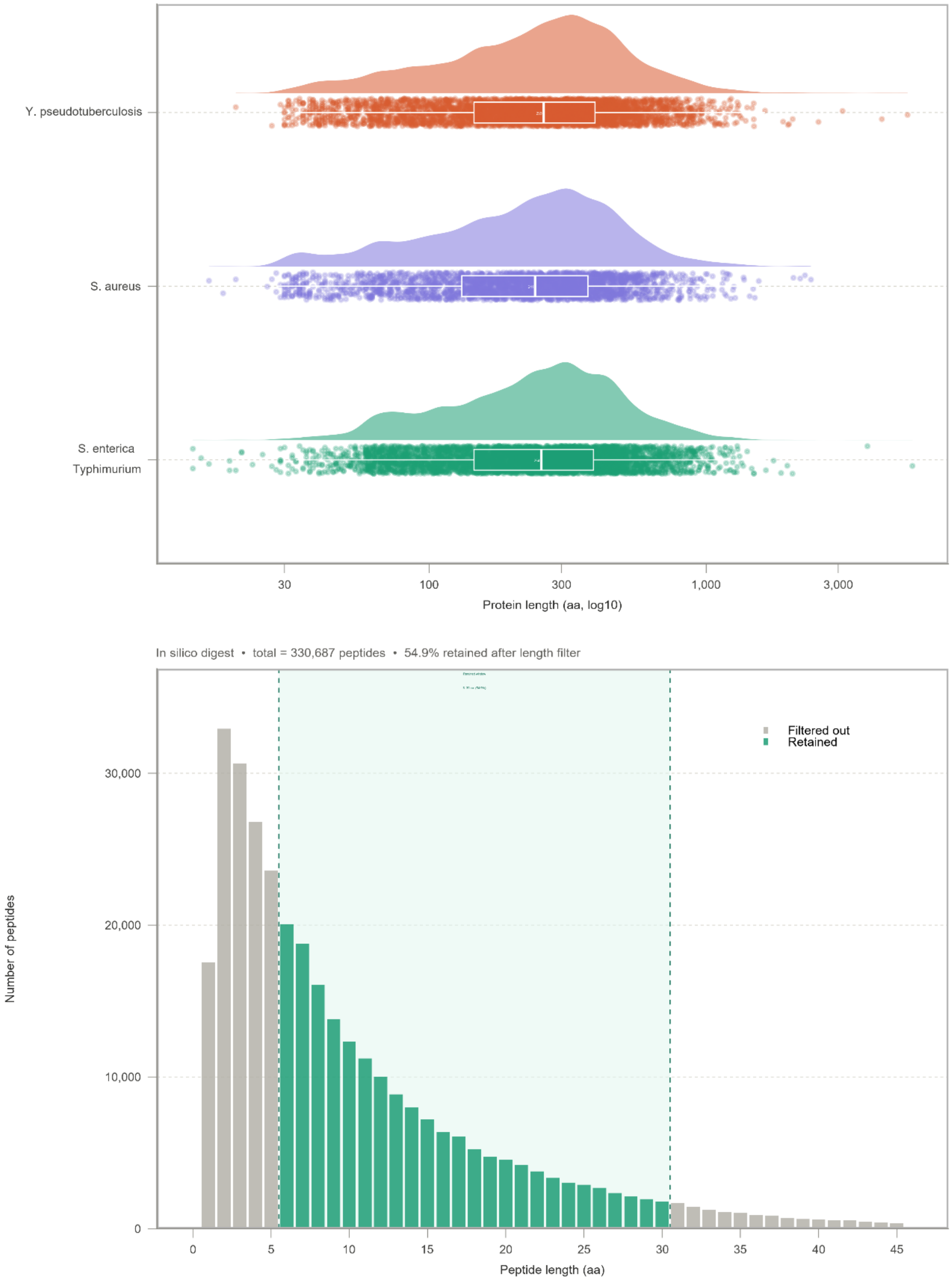


**Fig. S10. Protein and peptide length vary substantially within and across species.** A) Distribution of protein lengths across the three bacterial pathogens. Each point represents an individual protein, plotted on a log10-scaled x-axis. The shaded cloud reflects the density of proteins at each length, and the boxplot displays the median along with the first and third quartiles. B) Distribution of peptide lengths generated by trypsin digestion. Bars represent the density of peptides at each length. Peptides highlighted in green are those retained for iBAQ quantification, corresponding to tryptic peptides that fall within the expected detection range.

**Table**

**Tab. S1. iBAQ values for all genes detected by mass spectrometry for each replicate of 10 stress conditions and unexposed control conditions. The values for each species are shown in a separate sheet.**

**Tab S2. Differential expression levels of discordantly regulated genes in gene clusters with different expression patterns. Gene lists for each species are shown in separate sheets.**

**Tab S3. GO enrichment analysis on gene clusters with discordantly regulated mRNA and protein levels across three species. Results for each species are shown in separate sheets.**
